# Responses of dissolved organic matter to temperature change in the global ocean

**DOI:** 10.1101/2024.09.06.611638

**Authors:** Ang Hu, Yifan Cui, Sarah Bercovici, Andrew J. Tanentzap, Jay T. Lennon, Xiaopei Lin, Yuanhe Yang, Yongqin Liu, Helena Osterholz, Hailiang Dong, Yahai Lu, Nianzhi Jiao, Jianjun Wang

## Abstract

How dissolved organic matter (DOM) responds to climate warming is critical for understanding its role in future ocean carbon cycling. Here, we use a highly resolved dataset of over 800 DOM samples covering the surface waters to the deep Atlantic, Southern, and Pacific oceans to examine the changes in DOM molecular composition in response to warming water temperatures, referred to as DOM thermal responses. Towards the deep waters, the strength (i.e., overall magnitude) and diversity (i.e., variation among molecules) of these thermal responses both decline. However, these responses show opposite trends with the concentration of more recalcitrant molecules, decreasing and increasing, respectively. These contrasting trends concur with the observation that, compared to the strength of thermal responses, their diversity is more strongly explained by photochemical processes of DOM. By projecting global ocean thermal responses from 1950 to 2020 using environmental temperature, salinity and radiation, we predict that increases in the diversity of thermal responses are unexpectedly largest at deeper depths (> 1,000 m). Such increases could elevate the recalcitrant deep-ocean carbon sink by approximately 10 Tg C yr^-1^, which accounts for > 5% of the carbon flux reaching and persisting in the deep ocean. Our findings highlight the role of photochemical processes in imprinting DOM thermal responses, offering new insights into the future capacity of the oceanic carbon sink under global climate change.

---

Dissolved organic matter (DOM) in the ocean is one of the largest carbon reservoirs on Earth, holding ∼660 petagrams of carbon, which is more than marine and terrestrial biota combined ^1, 2^. Marine DOM responses to anthropogenic warming are crucial for understanding global carbon stocks and Earth’s climate ^3, 4, 5, 6^. However, predicting how, and to what extent, marine DOM responds to climate change is challenging. Past studies have primarily focused on coarse-level measurements such as ecosystem-scale carbon fluxes ^7, 8^, and the metabolic activities of microorganisms such as growth and respiration rates ^9, 10^, partially because the molecular composition of DOM is challenging to assess. DOM consists of thousands of individual organic molecules, each of which can respond differently to warming owing to their varying chemical properties ^11^. Bulk measurements of carbon may mask the influence of distinct molecules within a DOM sample (i.e., a DOM assemblage), overlooking the variability among these molecules, and be unable to differentiate their contributions to carbon persistence. Therefore, resolving how individual molecules respond to warming is necessary to identify the molecular mechanisms underlying changes in DOM composition and enable more accurate predictions of the fate of DOM.

A recent advance in forecasting ecosystem-level carbon dynamics is the development of an indicator to quantify the changes in DOM molecular composition in response to warming ^11^. This indicator, termed the “thermal response”, measures how the relative abundance of an individual DOM molecule changes along temperature gradients. Molecules with positive thermal responses become more abundant under warmer conditions, whereas those with negative responses decrease with warming. This indicator captures the collective temperature responses of individual DOM molecules and has revealed compositional change that helps anticipate the responses of carbon fluxes to future warming, such as applications to solid-phase extracted DOM (e.g., in lake ecosystems ^11^). For instance, the dominance of aliphatic and peptide-like molecules with positive thermal responses largely increases total carbon concentrations in lake sediments under warmer temperatures ^11^.

Overall, the indicator has a ‘strength’ dimension that quantifies the balance between negative and positive thermal responses among molecules within a DOM assemblage, describing the net magnitude of the dominant molecules’ changes with warming (Fig. 1). We propose that the indicator also has a ‘diversity’ dimension which describes the heterogeneity of these molecular responses across molecules (Fig. 1). This ‘diversity’ dimension captures diverse molecular properties reflecting complementary substrates available to decomposers or abiotic processing, which together can influence the transformation and fluxes of organic carbon ^12, 13^. Notably, the novel ‘diversity’ dimension of thermal responses and its influence on the DOM inventory, however, have been unexplored in previous studies.

**Figure 1.**
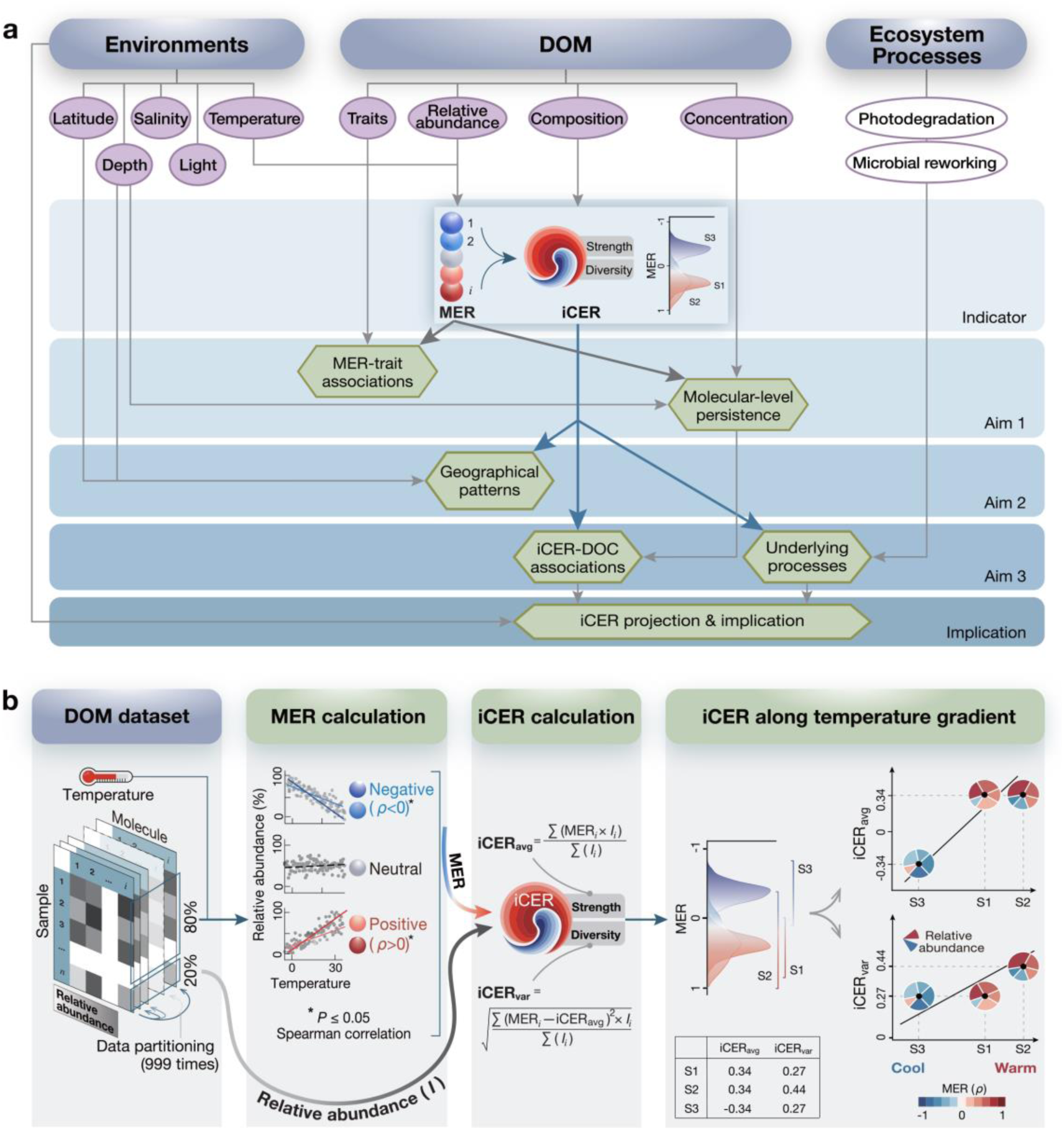
Study aims and indicator development workflow. (a) **Overview of study aims**. This study focuses on the indicator of compositional-level thermal response of DOM (termed “iCER”), quantified by aggregating the thermal responses of individual molecules (termed “MERs”). In the schematic, the Yinyang symbol illustrates sample-level iCER, with its Yin and Yang sections denoting the molecules with negative versus positive MERs. iCER has two dimensions: strength, representing the overall net response of individual molecules to temperature change, and diversity, representing the variation of these molecular responses among molecules. The development procedures of the indicators are shown in (b). We addressed three main aims: i) Aim 1 investigates the associations between MERs and molecular traits, and the molecular-level persistence of DOM inferred from non-significant depth profile of molecule-specific carbon concentrations. ii) Aim 2 explores changes in iCERs along latitude, water depth, and temperature. iii) Aim 3 examines the associations between iCERs and concentrations of dissolved organic carbon (DOC) or individual molecules, along with the underlying ecosystem processes, such as microbial reworking and photochemical degradation. iv) We predict spatiotemporal changes in iCERs in the global ocean and their implications for carbon sink dynamics. (b) **Workflow for indicator development**. The dataset of DOM molecular composition (relative abundances across all samples) was randomly partitioned into two independent subsets containing 80% and 20% of the samples, respectively, ensuring that each iCER was calculated using MERs derived from an independent subset of samples. MER was calculated as Spearman’s correlation coefficient *ρ* between the relative abundance of each molecule and corresponding temperatures. Molecules with negative, neutral, and positive MERs are represented by blue, grey, and red circles, respectively, with dark shading indicating higher magnitudes of MERs. Based on MERs, iCER strength (iCER*_avg_*) and diversity (iCER*_var_*) were calculated for each sample. To illustrate the meaning of these metrics, MER distributions were visualized using density plots, and the DOM compositional reorganization under warming was illustrated using pie charts for hypothetical three samples (S1–S3), each containing five molecules. The color gradient from blue to red in the ‘iCER’ circles and the density plots represents molecules with MERs ranging from negative to positive. Pie sections of a circle indicate the relative abundance of molecules with positive (red) and negative (blue) MERs.

The strength and diversity of thermal responses may be differentially driven by ecosystem processes affecting DOM turnover such as microbial and photochemical transformations. For example, microbial processing of DOM depends more deterministically on environmental conditions, such as higher temperatures that increase kinetic energy, enzymatic activities and primary productivity ^14, 15, 16, 17^. Additionally, enzymatic substrate specificity can increase the efficiency of microbial transformation of DOM by reducing the energetic and resource demands to maintain metabolic processes ^17, 18^. DOM assemblages structured by microbial processing are therefore expected to be dominated by molecules with strong positive thermal responses at higher temperatures, that is, increased strength of thermal responses. Although we focused on temperature-associated responses, we acknowledge that nutrient availability and other environmental factors can also regulate microbial DOM processing and may interact with temperature effects. In contrast, compared to the temperature-dependent and selective nature of microbial processes, abiotically-driven photochemical transformations of DOM are more stochastic and less constrained by environmental fluctuations, such as temperature. This is because photodegradation can proceed non-selectively across a wider range of molecules through highly reactive oxidizing agents, such as reactive oxygen species and radical intermediates ^19, 20^, which are primarily photon-driven rather than thermally activated. DOM assemblages shaped by photochemical processes are therefore expected to be composed of molecules with greater variability in thermal responses, that is a higher diversity of thermal responses.

Here, we used a global ocean molecular dataset ^21, 22, 23^, measured with Fourier transform ion cyclotron resonance mass spectrometry (FT-ICR MS), to examine the strength and diversity of thermal responses in DOM and the underlying ecosystem processes (Fig. 1a). The dataset comprises 821 samples at 124 stations from the entire water column across a 5,800 m depth gradient in the Atlantic, Southern, and Pacific oceans (Fig. S1). The thermal responses were examined along natural temperature gradients using a space-for-time substitution approach, which is widely applied to infer climate change responses of organisms ^24, 25, 26^ and organic carbon ^27^. For each sample, the strength and diversity of thermal responses were quantified based on each molecular formula’s (hereafter molecule’s) thermal response, termed a molecule-specific environmental response of temperature (MER) (Fig. 1b). The MER is defined by the magnitude (i.e., size) and direction (i.e., negative or positive) of the Spearman’s rank correlation coefficient between the relative abundance of each DOM molecule and water temperature in the corresponding sample. Negative and positive MERs correspond to DOM molecules that deplete or accumulate, respectively, as temperatures increase (i.e., warm-depleting and warm-accumulating molecules). More negative and positive MER values reflect greater net changes in a molecule’s relative abundance along the temperature gradient, indicating higher susceptibility of DOM molecules to temperature change. It should be noted that MER of a molecule refers specifically to its changes in relative abundance, but not absolute abundance, and is shaped by integrated biogeochemical processes, rather than a direct temperature effect on reaction kinetics. Then, the abundance-weighted average of MERs in each DOM assemblage was used to derive an indicator of the strength of thermal responses, termed the average of compositional-level environmental responses of temperature (iCER*_avg_*) ^11^. iCER*_avg_* estimates the collective net response of individual molecules to temperature change within a DOM assemblage at a given depth or latitude. More negative and positive iCER*_avg_* values suggest that a DOM assemblage is increasingly composed of molecules with greater susceptibility to temperature changes, as reflected by the dominance of warm-depleting and warm-accumulating molecules, respectively. To measure the diversity of thermal responses, we calculated the abundance-weighted standard deviation of MERs in each DOM assemblage, termed iCER*_var_*. iCER*_var_* estimates the extent of variation in individual molecules’ responses to temperature change relative to that of the entire DOM assemblage. Low values of iCER*_var_* indicate a DOM assemblage composed of molecules with similar MERs (that is, similar magnitudes of their susceptibility to temperature change), while high values indicate a DOM assemblage composed of molecules with divergent MERs.

With the new approaches, we addressed three main questions (Fig. 1a, Table S1): (1) How do the molecular-level thermal responses of DOM vary among individual molecules and how are they associated with chemical properties? (2) How do the compositional-level thermal responses of DOM (i.e., iCER*_avg_* and iCER*_var_*) vary along environmental and geographical gradients (latitude, water depth, and temperature)? and (3) How are these responses influenced by microbial versus photochemical processes, as quantified by the indices based on experimentally validated molecular markers ^23^? We then predict the spatiotemporal changes in iCER*_avg_* and iCER*_var_* in the global ocean and discuss how these responses might influence standing stocks of dissolved organic carbon (DOC) that play an important role in regulating the global climate (Fig. 1a, Table S1).

## Results and discussion

### Molecular-level thermal responses of DOM

In the Atlantic, Southern, and Pacific oceans, DOM thermal responses varied substantially among individual molecules (Fig. 2a). Most molecules were significantly (*P* ≤ 0.05) correlated in relative abundance with water temperature, with 38% (*n* = 1,610) showing negative responses and 36% (*n* = 1,541) showing positive responses (Fig. 2a). Across molecules, MER ranged from −0.72 to 0.83, with median values of −0.23 and 0.21 for negative and positive responses, respectively (Fig. 2a).

**Figure 2.**
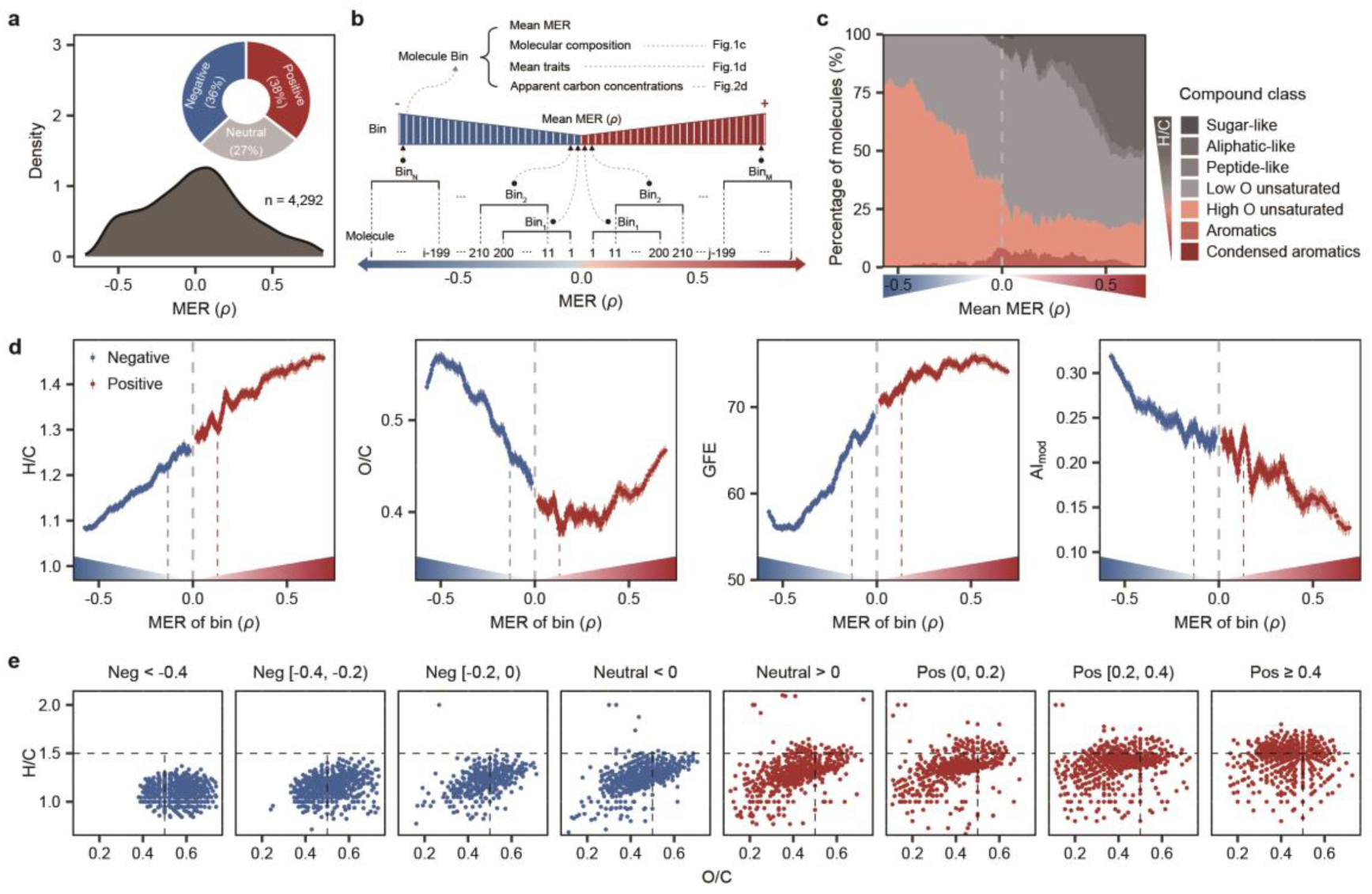
Molecule-specific thermal responses (MER) of DOM and their associations with molecular traits. (a) The distribution of MERs for DOM molecules across the global ocean. *n* is the number of molecules with MERs. Inset shows the percentage of molecules with negative, positive, and neutral MERs. (b) Cartoon illustrates a continuum of negative (blue) to positive (red) MERs. To visualize patterns along the MER gradient, all molecules were ordered by their MER values and grouped into 390 equal-sized and overlapping bins. Each molecule bin represents a DOM sub-assemblage, containing 200 molecules, with each successive bin offset by 10 molecules along the MER gradient. We further plotted (c) the molecular composition at the compound class level (i.e., the percentage of molecules within each compound class) and (d) means ± s.e of molecular traits along the MER continuum of the 390 bins. In (c), grey to red areas represent the compound classes with decreasing H/C ratios. In (d), we considered molecular traits, namely, H/C, O/C, Gibbs free energy (GFE), and the modified aromaticity index (AI_mod_). Blue and red dashed lines highlight the cutoffs of 100% significantly negative and positive MERs for the molecules in each bin, respectively. Lighter shaded lines denote the standard errors of molecular traits in each bin. (e) To visualize further H/C and O/C traits, we grouped all molecules into eight MER intervals, spanning a continuum from negative (Neg; blue) to positive (Pos; red) MERs, and displayed their distribution in van Krevelen diagrams. The MER ranges for each interval are shown at the top of panel (e).

The disparate thermal responses among individual DOM molecules could be explained in part by chemical traits, such as their recalcitrance and oxidation state (Figs. 2b-e). As MER became more positive along the continuum of thermal responses, over 75% of warm-accumulating molecules were classed as aliphatic-like and highly unsaturated compounds with low oxygen content in a DOM sub-assemblage, i.e. a bin of molecules classified by their MERs (see Methods; Figs. 2b, 2c). These molecules exhibited lower recalcitrance ^28, 29^, as indicated by an elevated H/C ratio and a lower aromaticity index (Figs. 2d, 2e). In contrast, as MER became more negative, warm-depleting molecules shifted towards highly unsaturated compounds with high oxygen content and higher recalcitrance, as indicated by a reduced H/C ratio from a mean of 1.25 to 1.08 and an increased aromaticity index from 0.23 to 0.32 (Figs. 2c-e). Their Gibbs free energy of oxidation also decreased from a mean of 69 to 58 kJ (mol C)^-1^ (Figs. 2d, 2e) indicating that these molecules with more negative MERs are thermodynamically less energy-rich and therefore provide less available energy for further microbial oxidation ^30^. Consistent with their elevated O/C and reduced H/C ratios (Figs. 2d, 2e), these thermodynamic shifts indicate that molecules with increasingly negative MERs fall towards the more recalcitrant end of the DOM reactivity spectrum.

We further assessed the persistence of these DOM molecules with negative or positive MERs throughout the water column by examining their apparent carbon concentrations (Fig. 3a), quantified with DOC-normalized peak intensities ^22, 31^. DOC-normalized peak intensities offer a semi-quantitative way to compare the abundance of the same molecular formula across samples ^32^ (see Methods eq. 3). We found that the recalcitrant molecules with negative thermal responses were evenly distributed in concentration throughout the water column, suggesting they persisted across the global ocean (Figs. 3, S2). For instance, the apparent carbon concentrations of individual molecules with negative MERs generally showed statistically non-significant changes with water depth, on average, at 82% of stations (*P* > 0.05; Figs. 3b, 3c). Similarly, as MER became more negative, fewer changes throughout the water column were observed in the composition and concentrations in each DOM sub-assemblage, i.e. MER-based molecule bin (Figs. 3d, S2). However, for individual molecules or the bins with more positive MERs, their apparent carbon concentrations decreased towards the deep ocean (Figs. 3c, S2). This pattern of the molecules with negative MERs aligns with the persistence concept of refractory DOC, which shows relatively persistent concentrations across water depths and may serve as an oceanic carbon sink ^33, 34, 35^.

**Figure 3.**
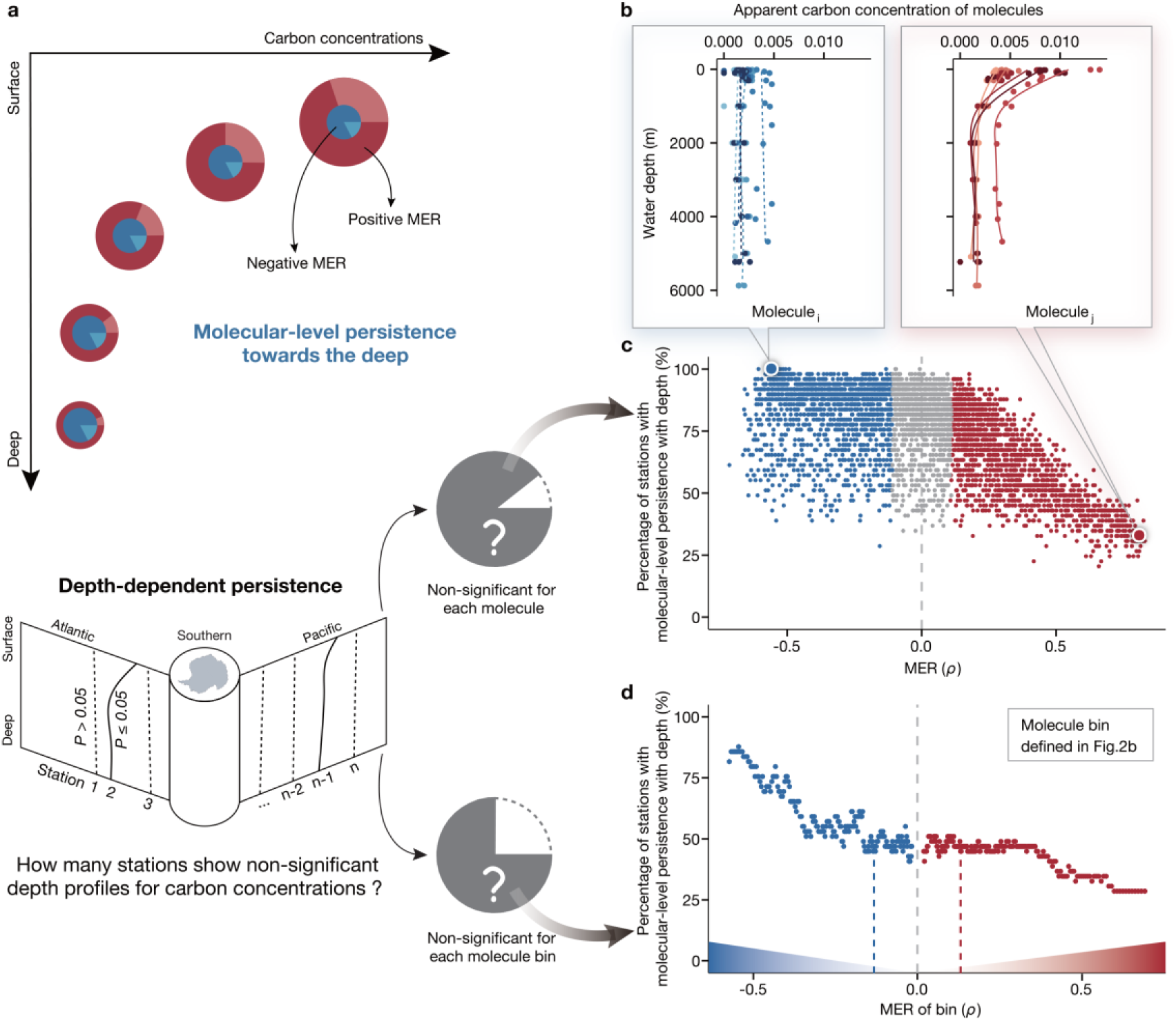
Depth profiles of carbon concentrations for DOM molecules along the MER continuum. (a) Conceptual illustration of how carbon concentrations may change for individual molecules with negative (blue) or positive (red) MERs from the surface to the deep ocean. In each pie, the size indicates bulk DOC concentration, and the four sections indicate the carbon concentrations of molecules with different MERs. At each station, molecular-level persistence could be reflected by statistically non-significant depth profiles of apparent carbon concentrations of DOM molecules. (b) Five stations are selected to illustrate the empirical depth profiles for two example molecules with negative (*m/z* = 391.06707, MER = −0.56; left panel) and positive (*m/z* = 317.1241825, MER = 0.81; right panel) MERs. Statistical significance of the depth profiles was tested with the two-sided Spearman’s rank correlation. Curves were estimated with generalized additive models with k = 5, where dotted curves indicate non-significant profiles (*P* > 0.05). (c-d) Across stations, the percentages of non-significant depth profiles are shown along the MER continuum. The depth profiles were considered for the apparent carbon concentrations of (c) each molecule individually and (d) each DOM sub-assemblage (i.e., the summed concentrations of molecules in each of the 390 molecule bins as in Fig. 2b). To clarify the calculation of percentages (i.e., the Y-axes), the procedures are illustrated in the left cartoon. For each molecule or molecule bin, the significance of depth profile was tested at a given station, and the percentage of non-significant depth profiles was then aggregated across all stations, represented by the grey portion of the pie. The larger the percentage of non-significant depth profiles, the longer the persistence of a molecule. Grey dots denote molecules with neutral MERs, while blue and red dots or triangles indicate negative and positive MERs, respectively. Blue and red dashed lines in (d) highlight the cutoff of 100% significantly negative and positive MERs for molecules in each bin, respectively.

### DOM thermal responses across latitudes and water depths

We quantified the strength (iCER*_avg_*) and diversity (iCER*_var_*) of thermal responses for each DOM assemblage by aggregating individual molecules’ MERs, and examined their geographical patterns. Across all samples, iCER*_avg_* ranged from −0.11 to 0.11, with a mean of −0.03 (Figs. 4a, 4b). Notably, 74% of DOM assemblages exhibited negative iCER*_avg_* values, indicating that molecules declining in abundance with increasing temperature dominate most ocean samples (Figs. 4a, 4b). iCER*_avg_* further significantly decreased towards higher absolute latitudes and greater water depths (*R*^2^ = 0.21 and 0.22, respectively; *P* < 0.001; Figs. 4a, 4b). The associations between iCER*_avg_* and absolute latitude varied across different water depths (Figs. 4a, S3). The slopes of linear regressions were negative (*P* ≤ 0.05) at shallower depths (≤ 200 m) but not significant (*P* > 0.05) at deeper depths (> 200 m) (Fig. S3). These results indicate that DOM assemblages at lower-latitude surface depths and at higher latitudes or deeper depths are increasingly composed of molecules with greater susceptibility to temperature changes, reorganizing into assemblages with higher abundances of warm-accumulating and warm-depleting molecules, respectively.

**Figure 4.**
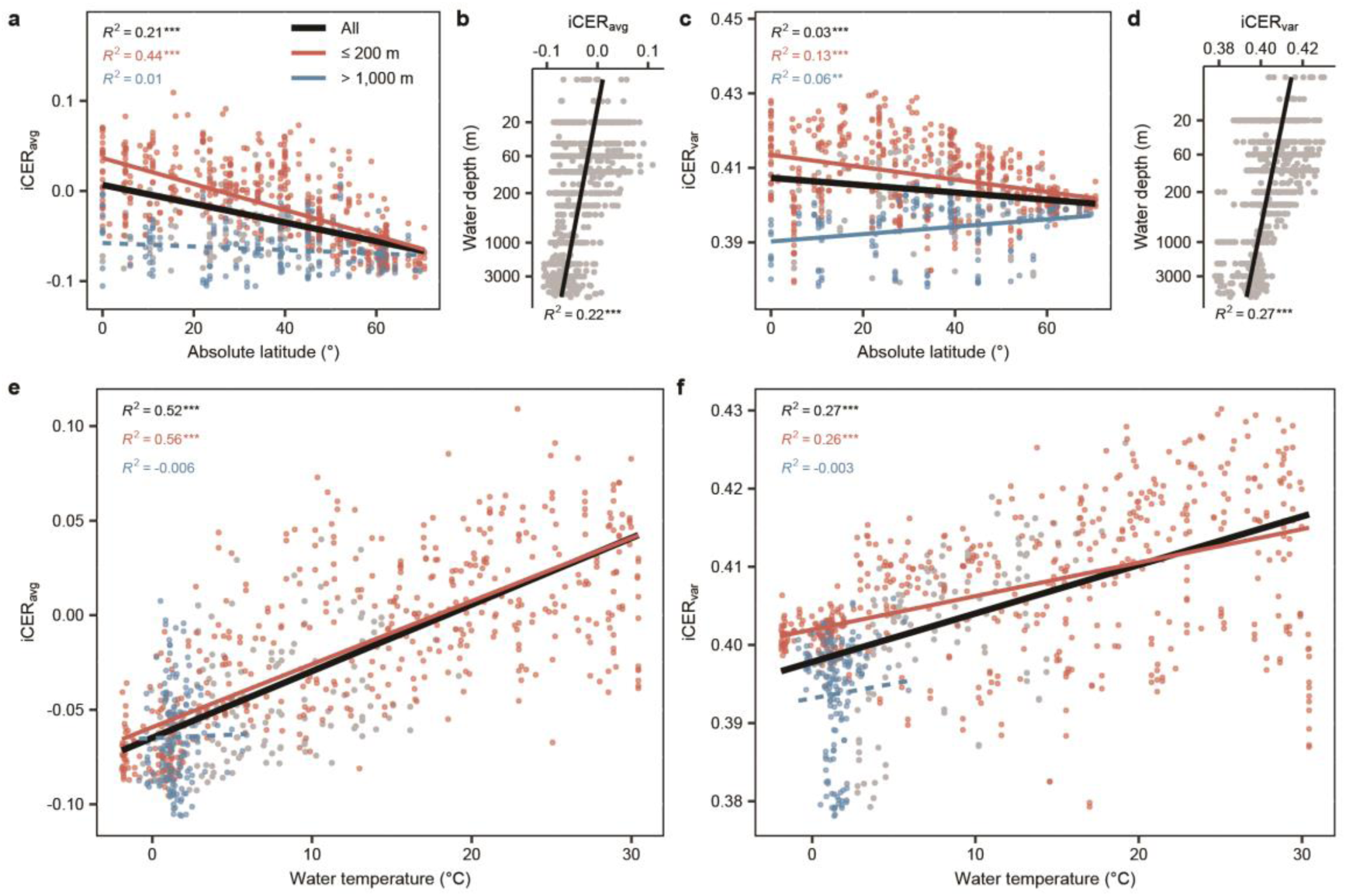
The compositional-level thermal responses of DOM along geographic and temperature gradients. We plotted the strength (iCER*_avg_*; a, b, e) and diversity (iCER*_var_*; c, d, f) of thermal responses in DOM assemblages against absolute latitude (a, c), water depth (b, d), and water temperature (e, f) for all samples (black lines, *n* = 821). For latitude and temperature, we considered samples in two depth categories of ≤ 200 m (red lines, *n* = 523) and > 1,000 m (blue lines, *n* = 159). Statistical significance of linear model fits is indicated by solid (*P* ≤ 0.05) or dotted (*P* > 0.05) lines. Only weighted linear regressions are shown for the iCER-temperature relationships, with statistical significance indicated by solid (*P* ≤ 0.05) or dotted (*P* > 0.05) lines.

Compared to the iCER*_avg_*, the diversity metric iCER*_var_* showed weaker latitudinal patterns but similarly strong depth-related patterns (Figs. 4c, 4d, S3). For instance, iCER*_var_* decreased towards high absolute latitudes with a lower *R*^2^ of 0.03, compared with an *R*^2^ of 0.21 for iCER*_avg_* (Figs. 4a, 4c). Across water depths, the linear slopes between iCER*_var_* and absolute latitude were negative (*P* ≤ 0.05) at shallower depths (≤ 200 m) and positive (*P* ≤ 0.05) at depths greater than 1,000 m (Figs. 4c, S3). These results indicate that, at low latitudes, individual DOM molecules in surface depths vary more in their thermal responses than in deep waters, reflecting a compositional reorganization of DOM towards molecules with more divergent susceptibility to temperature change.

The observed geographical patterns suggest that temperature plays an important role in explaining both iCER*_avg_* and iCER*_var_*. This is supported by the observation that both iCER*_avg_* and iCER*_var_* showed positive relationships with water temperatures, with *R*^2^ of 0.52 and 0.27, respectively (*P* < 0.001; Figs. 4e, 4f). We further found that the slope of linear regression between iCER*_avg_* and temperature was highest in the 60-100 m water layers, increasing iCER*_avg_* of 0.0036 per 1 °C, while for iCER*_var_*, the slope was greatest in the 300-1,000 m layers, increasing by 0.0016 per 1 °C (*P* ≤ 0.05; Fig. S3). This result suggests that the highest sensitivity of thermal responses to warming occurs at shallower depths for the strength of thermal responses than that for the diversity of responses.

### Implications of DOM thermal responses and underlying ecosystem processes

We found that the diversity of thermal responses was a better predictor of recalcitrant DOC concentrations than the strength of responses, especially in the deep ocean. At shallower depths (≤ 200 m), both iCER*_avg_* and iCER*_var_* exhibited positive relationships with DOC concentration, as well as with the apparent carbon concentrations of less recalcitrant molecules (that is, DOM sub-assemblages with more positive MERs) (*P* < 0.05; Figs. 5a-d). However, at deeper depths (> 1,000 m), iCER*_avg_* and iCER*_var_* showed contrasting relationships with DOC concentration (Figs. 5a-d). Specifically, iCER*_avg_* showed a marginally negative relationship with DOC (*R*^2^ = 0.03; *P* = 0.025), while iCER*_var_* had a statistically positive relationship (*R*^2^ = 0.22; *P* < 0.001; Figs. 5a, 5c). These contrasting patterns are in line with decreasing and increasing trends of iCER*_avg_* and iCER*_var_*, respectively, with the apparent carbon concentrations of more recalcitrant molecules (that is, DOM sub-assemblages with more negative MERs) (Figs. 5b, 5d). For instance, as MER became more negative, the linear slopes between the apparent carbon concentrations of DOM sub-assemblages with negative MERs and iCER*_avg_* became increasingly negative, while for iCER*_var_*, the slopes became increasingly positive, with both trends being pronounced at depths of > 1,000 m (Figs. 5b, 5d). Collectively, these results indicate that more diverse responses to warming among molecules can result in higher concentrations of recalcitrant DOC.

**Figure 5.**
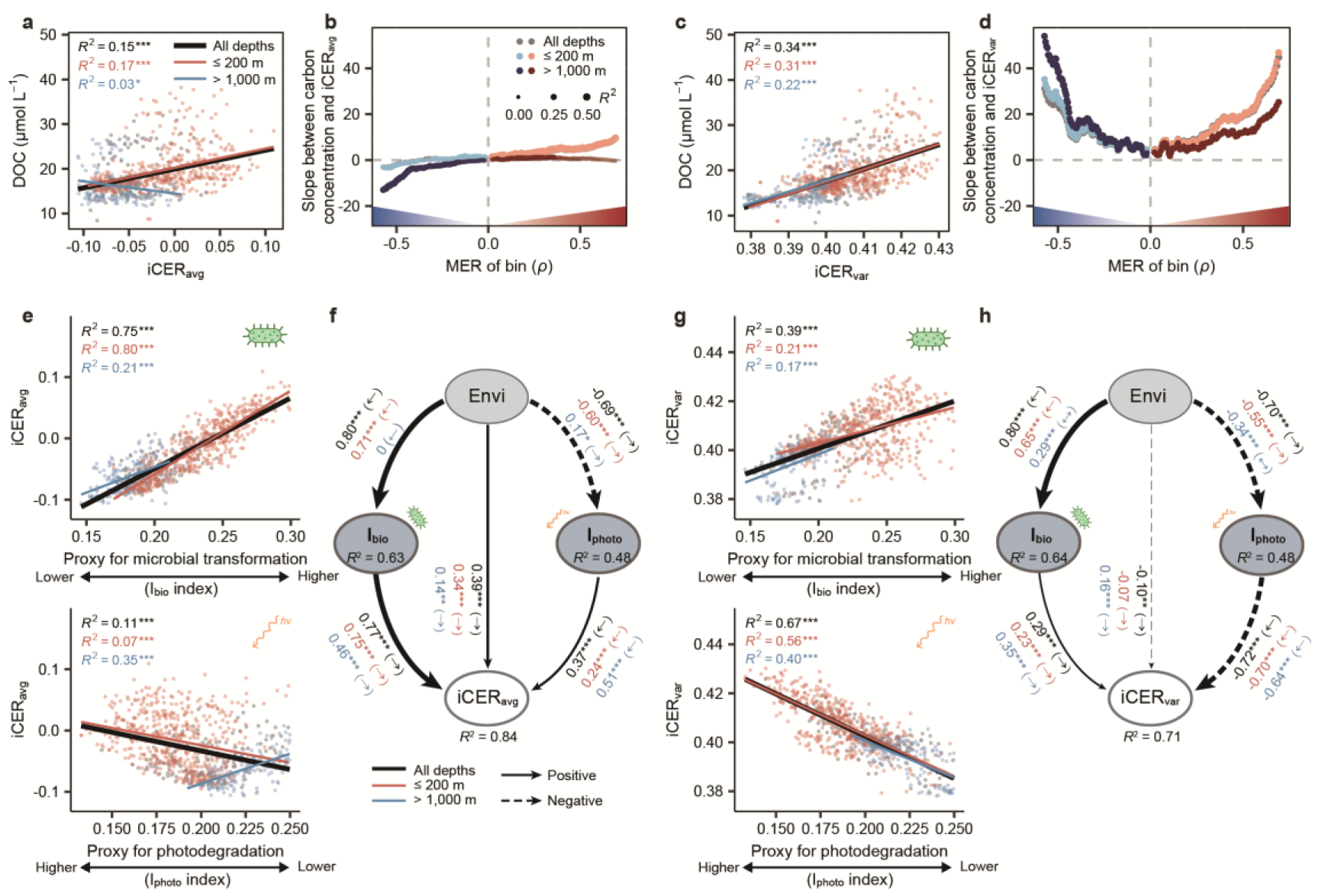
The compositional-level thermal responses of DOM linked to carbon concentrations and their underlying ecosystem processes. (a-d) **The relationships between iCERs and carbon concentrations**. We plotted SPE-DOC concentration against (a) iCER*_avg_* or (c) iCER*_var_*, and significant (*P* ≤ 0.05) linear model fits are indicated by solid lines. To further examine how the iCERs could predict recalcitrant DOC concentrations, we analyzed the linear regressions between (b) iCER*_avg_* or (d) iCER*_var_* and the apparent carbon concentrations of DOM sub-assemblage (i.e., the summed concentrations of molecules in each of the 390 molecule bins). The linear slopes were plotted along the MER continuum from negative to positive, corresponding to a gradient from more recalcitrant to more labile molecules. The adjusted *R*^2^ of regressions are indicated by dot sizes. (e-h) **The importance of microbial transformations (*I*_bio_) and photochemical degradation (*I*_photo_) in explaining variation in iCER*_avg_* and iCER*_var_*.** Changes in (e) iCER*_avg_* and (g) iCER*_var_* were plotted against *I*_bio_ or *I*_photo_, and significant (*P* ≤ 0.05) linear model fits are indicated by solid lines. Best-fitting structural equation models illustrate the indirect effects of environmental factors (Envi; details in Table S4) on (f) iCER*_avg_* and (h) iCER*_var_* mediated through *I*_bio_ and *I*_photo_. *R*^2^ denotes the proportion of variance explained for the endogenous variables. Dotted and solid arrows indicate negative and positive relationships, respectively. Arrow widths and accompanying numbers in black are the relative effects (that is, standardized path coefficients) of modeled relationships for all samples. The arrows in parentheses describe the modelled causal relationships, and asterisks represent significant effects at \*\*\**P* ≤ 0.001; \*\**P* ≤ 0.01; \**P* ≤ 0.05. Model fits are detailed in Table S5. The above analyses were also performed for the depth groups of ≤ 200 m and > 1,000 m. Numbers in red and blue (f, h) are the standardized path coefficients for the depths of ≤ 200 m and > 1,000 m, respectively.

The contrasting results between the strength and diversity of thermal responses could be explained by different ecosystem processes affecting DOM turnover (Figs. 5e-h). Microbial transformations and photochemical degradation are two key processes that control the production and consumption of DOM under specific environmental conditions such as temperature, salinity, and light ^2, 23, 36, 37^. The thermal responses of DOM are expected to be influenced directly by environmental factors and indirectly through the two ecosystem processes. For instance, environmental factors, such as water temperature, salinity, latitude, and water depth, explained 52-67% of the variations in both iCER*_avg_* and iCER*_var_*, except for iCER*_avg_* at deeper depths (> 1,000 m) with only 6% explained variation (Fig. S4). We then examined the effects of these two processes using the indices of *I*_bio_ and *I*_photo_, calculated for each sample. These two indices are based on the relative abundance of marker molecules that estimate the contributions of microbially reworked and photochemically degraded DOM, respectively ^23, 38, 39^. High *I*_bio_ and low *I*_photo_ values indicate that biological reworking and photodegradation, respectively, have shaped the current DOM composition ^23^.

Microbial transformations primarily explained the strength of thermal responses, particularly in surface waters. For instance, the iCER*_avg_* had a much stronger relationship with *I*_bio_ (*R*^2^ = 0.75; *P* < 0.001) than *I*_photo_ (*R*^2^ = 0.11; *P* < 0.001; Fig. 5e). When accounting for the effects of environmental factors such as water temperature and salinity using a structural equation model, *I*_bio_ still had more than twice the magnitude of direct effects on iCER*_avg_* (mean standardized effect = 0.77; *P* ≤ 0.001) compared to *I*_photo_ (mean effect = 0.37; *P* ≤ 0.001; Fig. 5f). At shallower depths, such effects of *I*_bio_ on iCER*_avg_* were more pronounced (*R*^2^ = 0.80; Figs. 5e, 5f). Consistent with the observed dominant effects of microbial transformations, iCER*_avg_* positively correlated with the DOM molecular lability indicated by elevated H/C ratio or Gibbs free energy of oxidation across both shallower and deeper depths (*R*^2^ = 0.88 to 0.94; *P* < 0.001; Fig. S5). These results collectively indicate that greater microbial production and transformation of DOM in warm and productive surface layers promotes the accumulation of more labile molecules as temperatures increase.

Interestingly, photochemical degradation was more important in driving the diversity of thermal responses than the strength across both surface and deep waters. Specifically, the iCER*_var_* depended more on photodegradation than microbial transformations. iCER*_var_* strongly declined with *I*_photo_ (*R*^2^ = 0.67; *P* < 0.001), but had a weaker relationship with *I*_bio_ (*R*^2^ = 0.39; *P* < 0.001), indicating a potentially more important role for photodegradation than microbial transformations (Fig. 5g). Even when accounting for the influences of environmental factors, the dominant effects of *I*_photo_ on iCER*_var_* remained, with *I*_photo_ having 2.5-times stronger direct effects compared to *I*_bio_ (mean standardized effect = −0.72; *P* ≤ 0.001 vs. mean effect = 0.29; *P* ≤ 0.001; Fig. 5h). The dominant effects of *I*_photo_ on iCER*_var_* were maintained across both shallower and deeper depths (Figs. 5g, 5h).

The above results indicate that photochemical processes could leave long-lasting influence (i.e., legacy effects) on deep-ocean DOM, mediated by environmental factors and physical mechanisms, thereby contributing to the elevated diversity of thermal responses. For instance, temperature showed a significant overall relationship with *I*_photo_ across all samples (*R*^2^ = 0.43; *P* < 0.001), which was also true at shallower depths (*R*^2^ = 0.37; *P* < 0.001; Fig. S6). This is consistent with the significant mean standardized effects of temperature on *I*_photo_ in the structural equation model (*P* < 0.001; Fig. 5h). In the euphotic zone, phytoplankton growth and DOC production are generally enhanced by warmer temperatures, potentially increasing the pool of DOM available for photochemical processing ^40, 41^.

Towards the deep ocean, the legacy of these surface photochemical processes is most likely explained by the export of partially photodegraded DOM, even though photodegradation itself generally occurs in the surface layer where solar radiation drives the generation of reactive oxygen species ^42, 43^. Nonselective photodegradation produces a wider range of DOM molecules with diverse molecular properties ^19^, and some partially photodegraded DOM, due to its relative recalcitrance, escapes microbial action upon export from the euphotic zone and persists in deeper waters ^23, 44, 45^. This process, combined with physical mechanisms, such as gravitational settling of particles and subsequent release of DOM as well as physical convective mixing, can contribute to transport of surface-derived organic matter to deeper waters ^1, 46, 47^. Consequently, the export of photodegraded DOM would elevate the diversity of thermal responses and the persistence of DOM in the deep ocean, reflecting the legacy effects of surface photochemical processes. In contrast, microbially transformed DOM produced in the surface waters may hardly reaches the deep ocean due to continuous remineralization during the long journey ^23, 48^. This explanation is supported by the molecular traits of deep-water DOM, which show increased molecular recalcitrance, indicated by a reduced H/C ratio or an elevated aromaticity index (*R*^2^ = 0.11 to 0.28; *P* < 0.001; Fig. S5).

### Predicting DOM thermal responses and carbon sink changes in the global ocean

We found that environmental variables can predict the spatiotemporal changes in the strength and diversity of thermal responses across the global ocean (Fig. 6). We trained a machine learning model − random forest ^49^ − to identify the variables that best explained iCER*_avg_* and iCER*_var_*. We analyzed 13 environmental variables, including water depth, the mean and seasonality of ocean temperature, salinity, and radiation attributes (see Supplementary Methods). Ocean temperature was most important for predicting iCER*<u>_avg_</u>*, while other variables were relatively unimportant (Fig. 6a). For iCER*_var_*, however, the most important variable was ocean temperature, followed by salinity, water depth, and the seasonalities of temperature (Temp.var), salinity (Salinity.var), and surface sensible heat flux (SSHF.var) (Fig. 6a). Overall, the environmental variables explained 70% and 62% of the variation in iCER*_avg_* and iCER*_var_*, respectively (Fig. 6a).

**Figure 6.**
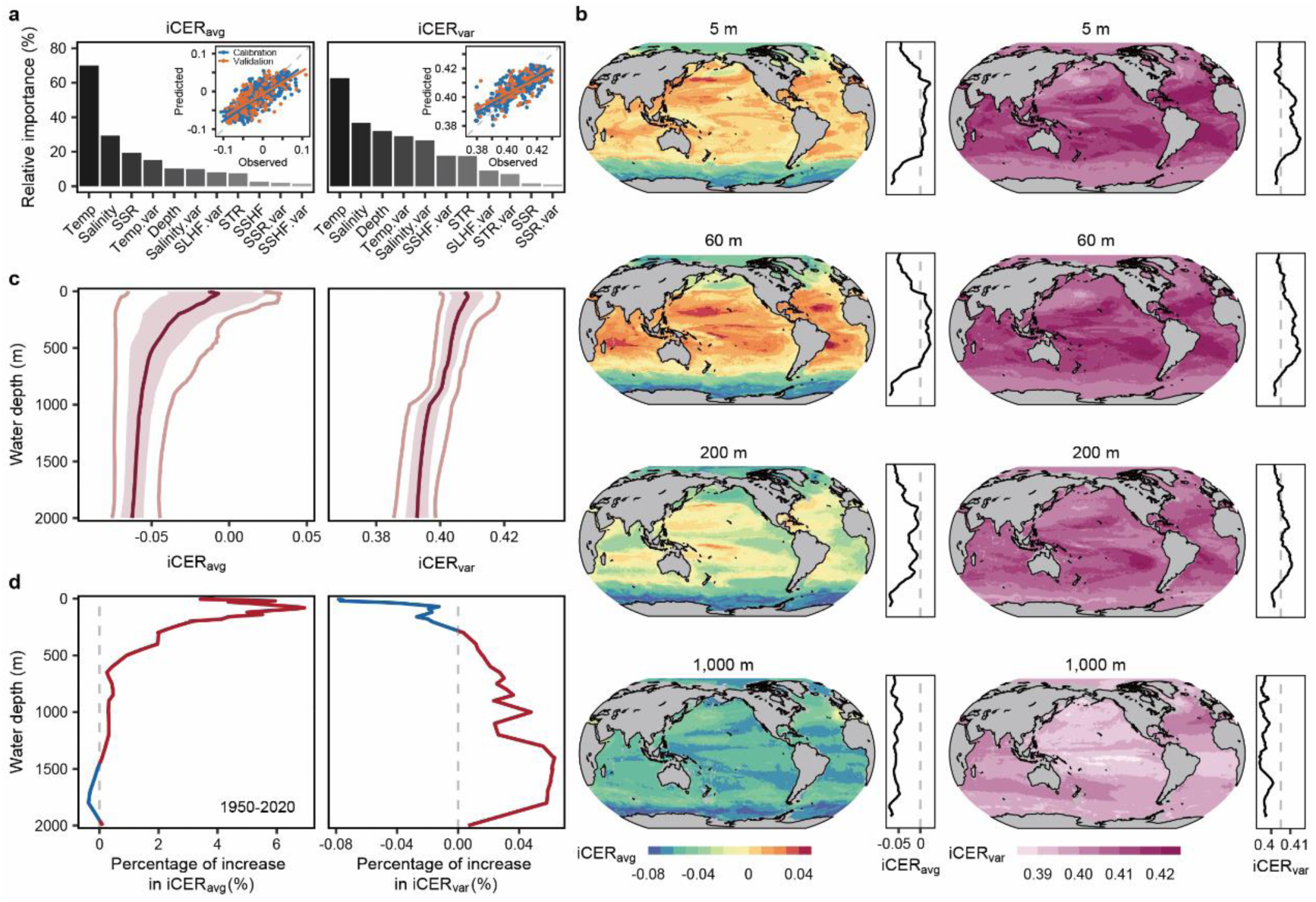
Thermal responses of DOM mapped across the global ocean and their changes since 1950. (a) The relative importance of environmental variables in influencing iCER*_avg_* and iCER*_var_* was estimated by random forest models. Insets show the model performance, with root mean square error (RMSE) and *R*^2^ for iCER*_avg_* (0.024 and 0.70, respectively) and for iCER*_var_* (0.006 and 0.62, respectively). Abbreviations of environmental variables see Supplementary Methods. Bar plots are used instead of distribution-based plots as variable importance is a model-derived summary statistic (b) Geographic distribution of iCER*_avg_* and iCER*_var_* in 2020. The four depths of 5, 60, 200, and 1,000 m represent the surface, deep chlorophyll maximum, mesopelagic, and bathypelagic waters, respectively. Global maps show modelled values at 1° spatial resolution and the adjacent panels show latitudinal trends in mean values. Vertical distributions of (e) the global mean iCER*_avg_* and iCER*_var_* in 2020 and (d) the percentage change in global mean iCER*_avg_* and iCER*_var_* between 1950 and 2020. Dark red line indicates global means and shading denotes their standard deviations. In (c), the light red lines on the left and right denote the lowest 5% and the highest 5% of global values at each latitude, respectively. In (d), blue and red lines indicate temporal decreases and increases, respectively.

The projected strength and diversity of thermal responses both declined towards the deep waters and showed little latitudinal variations in the deep ocean. In 2020, the global mean (±SD) thermal responses for iCER*_av<u>g</u>_* were −0.051 (±0.020) and 0.399 (±0.006) for iCER*_var_* (Figs. 6b, S7, S8). Towards greater water depths, the mean iCER*_avg_* and iCER*_var_* both declined from the surface until becoming practically invariant below ∼1,000 m, though the decline in iCER*_var_* was more gradual (Fig. 6c). At shallower depths (< 1,000 m), iCER*_avg_* and iCER*_var_* peaked in the subtropical oceans roughly between 20° and 40°, while at deeper depths (> 1,000 m), they were relatively low and more evenly distributed across latitudes (Figs. 6b, 6c, S7, S8). Both thermal responses were highest in subtropical shallower waters (< 1,000 m) likely because of high microbial and photochemical transformation and the export of surface-derived organic matter to the DOM pool in subtropical gyres ^16, 50, 51, 52^. These results are consistent with the observed geographical patterns (Fig. 4) and suggest that the DOM in subtropical shallower waters contains a higher abundance of warm-accumulating molecules compared to that in deeper waters or shallower waters at higher latitudes.

Since 1950, the deep ocean has generally experienced greater increases in the diversity of thermal responses compared to the strength. Specifically, the global mean (±SD) increases in iCER*_avg_* and iCER*_var_* from 1950 to 2020 were 0.922% (±1.622) and 0.028% (±0.028), respectively (Fig. 6d). Along the surface-to-deep-ocean gradient, iCER*_avg_* and iCER*_var_* exhibited significant temporal changes at over 95% of depths, most of which were positive at 72% and 85% of depths, respectively (Figs. 6d, S9). The largest temporal changes in iCER*_avg_* and iCER*_var_* occurred at different water depths. For iCER*_avg_*, the temporal changes were generally positive and large at shallower depths, with a maximum of 6.942% at 81 m, then sharply diminishing to zero and eventually becoming negative at deeper depths (> 1,457 m) (Fig. 6d). In contrast, the temporal changes in iCER*_var_* were negative only at shallower depths (0-283 m) but remained positive downwards, with a maximum of 0.063% at 1,401 m (Fig. 6d). Compared to iCER*_avg_*, the greater increases in iCER*_var_* in the deep ocean were further supported by the overall increase in their temporal changes. For instance, at depths of > 1,000 m, the temporal changes in iCER*_avg_* were almost negligible with a net increase of 0.005%, compared to a 5.649% increase at depths of < 100 m (Fig. 6d). However, the net increase in iCER*_var_* was substantially higher (0.047%) at depths of > 1,000 m than at depths of < 100 m (−0.039%) (Fig. 6d). Notably, the water temperature from 1950 to 2020 increased by an average of only 0.07 (±0.08) °C at deeper depths below 1,000 m versus 0.68 (±0.51) °C at 1 m depth (Fig. S10). Despite the relative stability of physical environments in the deep ocean, the substantial increase in the diversity of DOM thermal responses since 1950 indicates the changes in its molecular composition (specifically, greater heterogeneity in molecular thermal responses). These changes can be likely explained by the downward export of photochemically derived, recalcitrant molecules that imprint a persistent legacy on the deep-ocean carbon reservoir.

Our findings have important implications for the future of the deep-ocean carbon sink. This is largely because the diversity of thermal responses, rather than their strength, showed strong positive correlation with recalcitrant DOC especially at depths > 1,000 m (Figs. 5b, 5d). To illustrate the potential biogeochemical consequences, we estimated the apparent carbon concentrations of molecules with negative MERs (i.e., recalcitrant DOC) based on their relationship with iCER*_var_* (see Methods). Although peak intensities do not represent absolute molecular concentrations, they can provide reasonable proxies for relative abundances in the deep ocean, where the high molecular homogeneity of DOM composition markedly reduces variation in ionization efficiency. Using this semi-quantitative approach, the increase in iCER*_var_* from 1950 to 2020 corresponds to a potential rise of 9.87 (±3.35) Tg C yr^-1^ in the apparent concentration of recalcitrant DOC at depths > 1,000 m (Table S2). This magnitude is comparable to independent estimates of refractory DOC production rates (e.g., ∼43 Tg C yr^-1 53^) and thus represents a potentially important, previously unrecognized component of the long-term deep-ocean carbon budget. Given that ∼ 200 Tg C yr^-1^ of DOC is exported from the euphotic zone to depths > 500 m ^1^, our estimate implies that at least 5% of this flux could be influenced by changes in the diversity of DOM thermal responses. As the deep ocean stores about 72% of the global DOC inventory for years to centuries ^1^, future ocean warming is therefore likely to accelerate the accumulation of more persistent DOM at depth, effectively strengthening the role of the deep ocean as a long-term carbon sink.

It should be noted that these estimates reflect apparent rather than absolute concentrations and should be interpreted with appropriate caveats, as the FT-ICR-MS peak intensities are semi-quantitative due to differential ionization efficiencies among molecules. Nevertheless, for within-molecule comparisons across samples (that is, tracking the same molecular formula along environmental gradients), peak intensities can yield meaningful information on relative concentration changes, as supported by recent validation studies ^32^. This inference is particularly applicable when extended to subsets of molecules with negative MERs, which show remarkably low variations in apparent carbon concentrations across samples and depths (Fig. 3). Its reliability is further strengthened in the deep ocean (> 1,000 m), given that the DOM exhibits very high compositional homogenization (∼90% Bray-Curtis similarity on average ^54^). In this context, the large number of negative-MER peaks (> 1,500) could effectively average out molecule-specific ionization biases ^32^. Thus, matrix effects arising from interactions among co-occurring molecules during ionization are substantially reduced under conditions of compositional homogeneity and large ensemble size ^32^, which would improve the confidence of correspondence between summed peak intensities and true relative concentrations.

There are other uncertainties in the deep-ocean carbon sink, such as the dilution of low-concentration labile DOC ^55^, or the inputs of recalcitrant DOC from ancient deposits or hydrothermal circulation ^43, 56^, which were not considered here. Nevertheless, our estimates emphasize the pivotal but overlooked links between the diversity of thermal responses and both the magnitude and persistence of the global oceanic carbon sink. Future advances enabling direct quantification of molecule-specific concentrations will further refine these estimates and help reveal how global change reshapes the stability of the deep-ocean carbon pool.

We should also note that DOM composition is influenced by other environmental factors that may covary with temperature along oceanographic gradients, such as nutrients, light availability, and primary productivity. Consequently, MERs and iCERs capture how DOM molecular composition varies with *in situ* temperature while implicitly integrating these co-varying factors, rather than single-factor effects. This integration of multiple correlated factors is an inherent feature of space-for-time substitution approach, which captures main environmental gradients ^24^. Nonetheless, future work involving temperature-controlled experiments, direct measurements of reaction rates, and mechanistic models that incorporate multiple environmental drivers would be essential for disentangling the processes underlying these thermal responses.

## Conclusions

In conclusion, our study provides insights into how, and to what extent, oceanic DOM responds to climate warming by considering the strength and diversity of thermal responses among molecules. The findings indicate that the diversity of thermal responses is a better predictor of recalcitrant DOC concentrations than the strength. The deep ocean, conventionally considered a quasi-steady environment, is projected to have experienced an increasing diversity of thermal responses since 1950. This increase corresponds to an accumulation of approximately 10 Tg C yr^-1^ in recalcitrant deep-ocean carbon sink, mainly owing to the legacy effects of photochemical processes. Our results imply that the diversity of thermal responses in the deep ocean is sensitive to a warmer climate and could act as a negative feedback mechanism in the future, which needs to be taken into account in future models of the ocean carbon budgets ^57, 58^. We encourage future studies to provide experimental evidence on the effects of photochemical processes on the diversity of DOM thermal responses, and explore the interactions between photochemically-driven carbon sequestration and biological mechanisms, such as biological carbon pump ^6^. These studies would enhance our understanding of future ocean carbon storage under global climate change.

## Methods

### Ocean DOM composition data

To examine the DOM responses to temperature change, we used a global ocean DOM dataset spanning broad geographical gradients. This dataset, which is available from previous literature ^21, 22, 23^, was generated by the Research Group for Marine Geochemistry at the Institute for Chemistry and Biology of the Marine Environment (Carl von Ossietzky University Oldenburg, Germany). Briefly, the DOM dataset, measured with Fourier transform ion cyclotron resonance mass spectrometry (FT-ICR MS), comprises 821 samples collected from 124 stations across depth gradients ranging from 5 to 5,874 m during six research cruises in the Atlantic, Southern, and Pacific oceans (Fig. S1). Latitudinal transects were from 70.5°S to 43°N in the Atlantic and Southern Oceans and from 52°S to 59°N in the Pacific Ocean. All samples were collected from a rosette of Niskin samplers equipped with sensors for measuring water depth, temperature, and salinity. DOM of water samples was solid-phase extracted (SPE) ^59^ and measured using FT-ICR MS (solariX XR^TM^ system, Bruker Daltonik, Bremen, Germany) equipped with a 15 Tesla superconducting magnet and a standard electrospray ionization interface. Further details of sample collection, DOM extraction, and FT-ICR MS analysis of DOM composition are provided in previous literature ^21, 23, 60^ and briefly described in Supplementary Methods. We note that this study focuses on SPE-DOC (rather than bulk DOC) because it is the analytically attainable fraction for molecular DOM characterization via FT-ICR MS.

In total, 6,114 peaks had formula assignments for 821 samples and these formulae were referred to as “molecules” throughout the manuscript. The relative abundance (or intensity) of molecules was calculated by normalizing signal intensity of each peak to the sum of all assigned intensities within each sample. These molecules were categorized into seven compound classes based on van Krevelen diagrams ^61^ and modified aromaticity index (AI_mod_) ^62, 63^. The compound classes were defined as follows: low O unsaturated (AI_mod_ < 0.5, H/C < 1.5, O/C < 0.5), high O unsaturated (AI_mod_ < 0.5, H/C < 1.5, O/C = 0.5-0.9), aliphatic-like (H/C = 1.5-2.0, O/C < 0.9, N = 0), peptide-like (H/C = 1.5-2.0, O/C < 0.9, N > 0), sugar-like (H/C > 2.0 or O/C ≥ 0.9), aromatics (AI_mod_ = 0.5-0.67), and condensed aromatics (AI_mod_ ≥ 0.67). It should be noted that compounds identified as “aliphatic-like”, “peptide-like”, and “sugar-like” have the molecular formulae of aliphatics, peptides, and sugars, but their actual structure may differ ^63^. We evaluated four molecular traits, namely, H/C, O/C, standard Gibbs Free Energy (GFE) ^30^, and AI_mod_ ^62, 64^, which provide insight to the recalcitrance and oxidation state of DOM. DOM composition and characteristics are detailed in Supplementary Methods.

### Development of the indicator of compositional-level thermal responses

We developed the indicator to quantify how and to what extent DOM assemblages respond to temperature change (see Fig. 1a “Indicator” and Fig. 1b). The indicator, termed compositional-level environmental responses of temperature (hereafter “thermal responses”, abbreviated as “iCER”), is characterized by two key dimensions: the strength of these thermal responses (“iCER*_avg_*”) and their diversity (“iCER*_var_*”). iCER*_avg_* estimates the collective net response of individual molecules to temperature change within a DOM assemblage at a given depth or latitude, while iCER*_var_* estimates the extent of variation in individual molecules’ responses to temperature change relative to that of the entire DOM assemblage. The indicator development involved two primary procedures (Fig. 1b): (1) calculating each molecule’s thermal response, termed molecule-specific environmental response of temperature (MER), and (2) producing the metrics of iCER*_avg_* and iCER*_var_* for each sample. The inputs of our procedures are the relative abundance table of DOM composition and the corresponding water temperature across the 821 samples, and the outputs are the iCER*_avg_* and iCER*_var_* for each sample.

Our procedures begin by calculating MERs for individual DOM molecules across samples (Fig. 1b). The MER is defined as both the magnitude (i.e., size) and the direction (i.e., negative or positive) of the Spearman’s rank correlation coefficient (*ρ*) calculated between the relative abundance of each DOM molecule and the water temperature in the corresponding sample ^11^. To correct for the large number of tests performed, we applied a false discovery rate correction ^65, 66^, and correlations were considered statistically significant with an adjusted *P* ≤ 0.05. Negative, positive, and neutral MERs were defined as Spearman *ρ* < 0 (*P* ≤ 0.05), *ρ* > 0 (*P* ≤ 0.05), and other *ρ* (*P* > 0.05), respectively. Larger negative and positive Spearman *ρ* values indicate stronger negative and positive thermal responses of DOM molecules (MERs), respectively. Negative and positive MERs correspond to DOM molecules that deplete and accumulate, respectively, as temperatures increase (i.e., warm-depleting and warm-accumulating molecules). To reduce Type I errors in the correlation calculations created by low-occurrence molecules, we applied the majority rule, retaining only those molecules observed in more than 20% of the total samples.

The next step in the procedures calculates iCER*_avg_* and iCER*_var_* for each sample (Fig. 1b). We synthesized the MERs of all molecules within a DOM into two compositional-level metrics of iCER*_avg_* and iCER*_var_*. The iCER*_avg_* and iCER*_var_* were calculated as the relative abundance-weighted mean and standard deviation of MERs, respectively, for each sample, using the equations:

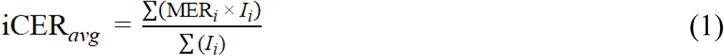

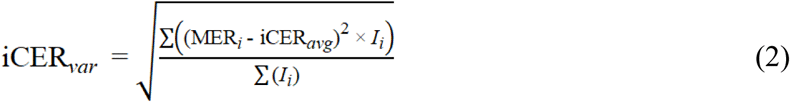

where MER*_i_* and *I*_i_ are the MER and relative abundance, respectively, for each individual molecule *i* within each sample. For the strength metric iCER*_avg_*, more negative and positive values suggest that a DOM assemblage is increasingly composed of molecules with greater susceptibility to temperature changes, as reflected by the dominance of warm-depleting and warm-accumulating molecules, respectively. An iCER*_avg_* of zero suggests that a DOM assemblage contains a mixture of molecules with both negative and positive MERs that can cancel each other out. For the diversity metric iCER*_var_*, low values indicate a DOM assemblage composed of molecules with similar MERs (that is, similar magnitudes of their susceptibility to temperature change), while high values indicate a DOM assemblage composed of molecules with divergent MERs.

By using the DOM dataset, we applied a space-for-time substitution approach ^26, 67^ to infer long-term responses of DOM composition to temperature change (i.e., “thermal response”) at both molecular and compositional levels from spatial environmental gradients. In this context, the long-term *in situ* processes structuring DOM under climate change can be inferred by comparing sampling sites with different temperatures along geographical gradients. This approach has been widely used in ecological studies to infer long-term responses from spatial environmental gradients, including applications to plants ^24, 25^, animals ^26, 67^ and organic matter ^27^. Such long-term responses could be rarely captured by short-term experiments, especially for marine DOM molecules, whose turnover times in the global ocean can reach 1,000 to 6,000 years ^68, 69^. Thus, our interpretation of MER and iCER relies on the assumption that long-term environmental regimes structure DOM composition, such that spatial gradients of temperature reflect persistent biogeochemical conditions. Under this assumption, temperature influences DOM composition indirectly by altering the relative contributions of multiple biogeochemical processes such as microbial transformations and photochemical degradation.

Practically, to ensure that the relative abundances of individual molecules used for iCER*_avg_* (or iCER*_var_*) calculations were independent from those used to derive MERs, we implemented a dataset-partitioning procedure prior to all indicator calculations. The entire input dataset was randomly divided into two independent subsets, containing 80% and 20% of the samples, respectively, as described in Hu *et al.* ^11^ (Fig. 1b). The dataset containing 80% of the samples was used to calculate the MERs of individual DOM molecules. Following the majority rule, molecules occurring in at least 20% of these samples (≥ 131 samples) were retained for MER calculation. The dataset containing the remaining 20% of the samples was then used to calculate iCER*_avg_* and iCER*_var_* for each sample, based on the relative abundance of individual molecules and their corresponding MERs. To obtain iCER*_avg_* and iCER*_var_* for all samples, the random partitioning and indicator-calculation procedures were repeated 999 times. For each sample, the final iCER*_avg_* and iCER*_var_* values were computed as the mean across all iterations for use in subsequent statistical analyses. Notably, MERs derived from all samples were nearly identical to those obtained by averaging the results of 999 random iterations using 80% of the samples, with a regression slope of 1 along the 1:1 line, indicating that MER calculations were highly stable and robust to data partitioning. After filtering out low-occurrence molecules (i.e., those present in less than 20% of the samples), 4,292 molecules were retained for subsequent molecular-level statistical analyses.

### Statistical analyses

#### (1) Molecule-specific thermal responses associated with molecular traits

We assessed how the molecular composition and traits (e.g., recalcitrance and oxidation state) of DOM changed towards stronger thermal responses of individual molecules (see Fig. 1a “Aim 1”; Table S1). To examine the continuous and/or sharp transitions in molecular composition or traits along thermal response gradient, we created a MER continuum by categorizing all molecules into equal-sized bins through a moving-window approach ^70^. Practically, we first separated the molecules into two groups with negative and positive MERs, and sorted the molecules of each group along MER gradients, that is, from the lowest to highest absolute values of MER. We categorized the molecules into 189 and 201 equal-sized bins for the groups with negative and positive MERs, respectively, according to the magnitude of MER values (Fig. 2b). Each molecule bin represents a DOM sub-assemblage consisting of 200 molecules that share similar magnitudes of thermal responses. Consecutive bins partially overlap with a step size of 10 molecules (i.e., 1–200, 11–210, …, 1881–2080 for groups with negative MERs, and 1–200, 11–210, …, 2001–2200 for groups with positive MERs). We then calculated the mean MERs for each of the 390 bins, resulting in a MER continuum ranging from −0.57 to 0.69. For each bin, we analyzed the molecular composition at the compound class level with respect to molecule occurrence, defined as the percentage of molecules within each compound class relative to the 200 assigned peaks, and calculated the mean (± s.e) values of individual molecular traits (including H/C, O/C, GFE, and AI_mod_).

#### (2) Depth profiles of apparent carbon concentrations of DOM molecules

We assessed the persistence of the DOM molecules with negative or positive MERs throughout the water column by analyzing their apparent carbon concentrations (see Fig. 1a “Aim 1”; Table S1). Persistence of DOM molecules was indicated by statistically non-significant changes in the apparent carbon concentrations of molecules with water depths at each station. The apparent carbon concentrations were quantified as DOC-normalized peak intensities, which are not absolute concentrations but provide robust within-peak comparisons across depths ^32^.

Specifically, we first quantified the apparent carbon concentrations for each molecule individually or each DOM sub-assemblage (i.e., the summed concentrations for the molecules in each of the 390 molecule bins mentioned above). The concentration for a molecule *i* (DOC_*i*_) was calculated as:

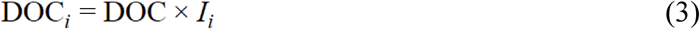

where DOC is the concentration of SPE-DOC (µM) and *I_i_* is the relative abundance of individual molecules. The concentration for a DOM sub-assemblage, i.e., multiple molecules in bin *j* (DOC_*j*_), was summed across these molecules:

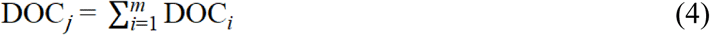

where *m* is the number of molecules in each bin.

At each station, we tested the statistical relationships between the apparent carbon concentrations of DOM molecules and the water depth using two-sided Spearman’s rank correlation. For each molecule bin, we tested the statistical relationships between its compositional dissimilarity (Bray-Curtis) and the difference in depth using Mantel test with 999 permutations. Across the stations, we then calculated the percentages of non-significant relationships described above for individual molecules or the molecule bins.

#### (3) Geographical patterns of compositional-level thermal responses

We analyzed the relationships of iCER*_avg_* or iCER*_var_* with latitude, water depth, and water temperature using linear models, with a F-statistic used to test the statistical significance of the regressions (see Fig. 1a “Aim 2”; Table S1). To evaluate heteroscedasticity, we additionally applied the Breusch-Pagan test ^71^, which identified non-constant variance in the iCER–temperature relationships. Accordingly, we re-fitted these models using weighted linear regression, which assigns lower weights to sample groups exhibiting higher residual variance (e.g., warm surface waters). Both modelling approaches produced consistent results and we thus reported only the weighted linear regression outputs in the figures. The latitudinal or temperature patterns of iCER*_avg_* and iCER*_var_* are likely little affected by DOM extraction efficiency or instrument conditions (see Supplementary Methods and Table S3).

To evaluate whether observed relationships between iCER*_avg_* or iCER*_var_* and environmental variables (e.g., temperature, latitude, depth) could arise by chance, we implemented a permutation-based null test in which temperature labels were randomly shuffled across samples (1,000 permutations). iCER*_avg_* and iCER*_var_* were recalculated under each permutation using the same 999-iteration procedure. The observed correlations between iCER*_a vg_* or iCER*_var_* and environmental variables (temperature, depth, latitude) lay far outside the distribution generated by the null models (*P* < 0.001), confirming that these relationships are unlikely to result from statistical artifacts or circularity.

To compare the results between the surface and deep ocean in the above analyses, we used representative depths of ≤ 200 m and > 1,000 m, respectively. The observed results are unlikely affected by variation in depth coverage based on the analyses across finer depth intervals (see Supplementary Methods). Furthermore, we provided additional support to compare the results between the surface and deep ocean based on water masses, and confirmed consistent results (see Supplementary Methods and Fig. S11).

#### (4) Linking compositional-level thermal responses to carbon concentrations and their underlying environmental factors and ecosystem processes

To examine how compositional-level thermal responses could predict recalcitrant DOC concentrations, we analyzed the relationships between iCER*_avg_* (or iCER*_var_*) and the apparent carbon concentrations of DOM sub-assemblages across a MER continuum from negative to positive, corresponding to a gradient from more recalcitrant to more labile molecules (see Fig. 1a “Aim 3”; Table S1). As in the previous analysis, the apparent carbon concentrations of DOM sub-assemblages across a MER continuum were calculated as the summed concentrations of molecules in each of the 390 molecule bins. Statistical relationships were tested using the adjusted *R*^2^ or the slope of linear models.

We further assessed the environmental factors and ecosystem processes underlying iCER*_avg_* and iCER*_var_* using random forest analysis ^72^, Pearson correlation, and structural equation modelling (SEM) ^73^ (see Fig. 1a “Aim 3”; Table S1). Environmental factors included water temperature, salinity, water depth, and latitude. We considered two key ecosystem processes—microbial transformations and photodegradation—which were quantified using the indices of *I*_bio_ and *I*_photo_, respectively ^23^ (see Supplementary Methods). SEM analysis can explore how environmental factors influenced iCER*_avg_* and iCER*_var_* directly and indirectly through the two ecosystem processes. More statistical methods are detailed in Supplementary Methods and Tables S4 and S5.

#### (5) Global mapping of compositional-level thermal responses

We estimated the spatiotemporal changes in iCER*_avg_* and iCER*_var_* across the global ocean in 1950 and 2020 by including 13 environmental variables in a machine learning-based random forest model (see Fig. 1a “Implication”; Table S1). Water depth and 12 other global environmental variables were extracted from global mapping products (see Supplementary Methods). These variables explained similar variation to the *in situ* measured variables (Fig. S12), but are more easily obtained for the historical and future modelling of global ocean.

We trained the random forest model to identify the most important variables explaining the variation in iCER*_avg_* and iCER*_var_*. The model with the lowest root mean square error (RMSE) was applied to predict iCER*_avg_* (or iCER*_var_*) in each 1° × 1° pixel across the global ocean. The mapping analysis was performed at 39 water depths for the upper 0–2,000 m in 1950 and 2020. We then examined the temporal changes in iCER*_avg_* and iCER*_var_* between 1950 and 2020 in each 1° × 1° pixel across the global ocean. The random forest model and the calculations of percentage changes are detailed in Supplementary Methods, Fig. S13, and Table S6.

We estimated the impact of changes in iCER*_var_* from 1950 to 2020 on the concentration changes of recalcitrant DOC in the global ocean (see Fig. 1a “Implication”; Table S1). The concentration of recalcitrant DOC was calculated as the apparent carbon concentrations of molecules with negative MERs (Spearman *ρ* < 0; *P* ≤ 0.05) and divided by DOM extraction efficiency. We used negative MERs because these molecules were more recalcitrant and persistent, as shown in Figs. 2 and 3. The estimate was based on the rate of change in the apparent carbon concentration of molecules with negative MERs with respect to iCER*_var_*:

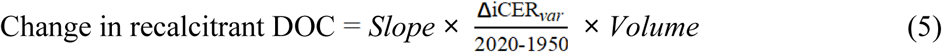

where *Slope* is the linear slope of the relationship between the iCER*_var_* and the apparent carbon concentration of molecules with negative MERs. ΔiCER_*var*_ is the change in iCER*_var_* from 1950 to 2020. *Volume* is the volume of seawater ^74^. The concentration of recalcitrant DOC was calculated separately for depths of < 100, 101-500, and > 1,000 m.

## Data availability

The DOM and associated environmental data in the global ocean are obtained via PANGAEA Data Publisher (DOI: 10.1594/PANGAEA.962747). Source data for all main figures are provided with this paper.

## Code availability

The main scripts and example data for DOM thermal response calculations are available at http://github.com/jianjunwang/iDOM.

## Acknowledgements

We acknowledge Cunjie Zhang, Shushi Peng, Gangsheng Wang, Xiancai Lu, Jizhong Zhou, and Guoping Zhao for support and comments. This study was supported by National Natural Science Foundation of China (42225708, 92251304, 42377122, U24A20578), Global Ocean Negative Carbon Emissions (Global ONCE) Program, Basic Research Program of Jiangsu (BK20240111), Key Laboratory of Lake and Watershed Science for Water Security (NKL2023-QN04), and Science and Technology Planning Project of NIGLAS (NIGLAS2022GS09).

