## Supplementary materials for "Responses of dissolved organic matter to temperature change in the global ocean"

**The file includes:**

Supplementary Methods

Supplementary Tables (S1–S6)

Supplementary Figures (S1–S13)

### Supplementary Methods

#### FT-ICR MS analysis of DOM sample

DOM of water samples was solid-phase extracted (SPE) for Fourier-transform ion cyclotron resonance mass spectrometry (FT-ICR MS) measurement [1]. Briefly, 4 L of seawater were filtered through a pre-combusted (400 ºC, 4 h) 0.7-µm glass fiber filters (GF/F, Whatman, United Kingdom), and the filtered water was acidified to pH 2 using 25% HCl and further extracted using 1 g PPL cartridges (Agilent Technologies). Cartridges were rinsed with pH 2 ultrapure water and dried with $N_2$ gas. The DOM was then eluted with 6 ml of methanol (HPLC-MS grade) into pre-combusted amber glass vials and immediately stored at -20 °C until FT-ICR MS analysis. The SPE-DOC concentrations were analyzed on a Shimadzu TOC-VPCH total organic carbon analyzer. The mean (±SD) carbon-based extraction efficiency was 53 ± 9% ($n$ = 285) for a representative subset of DOM samples, covering the entire water column and latitudes from 70°S to 43°N. Detailed methods for DOM extraction and extraction efficiency have been described previously [2, 3].

Highly accurate mass measurements of DOM extracts were conducted using FT-ICR MS. For FT-ICR MS analysis, the extracts were mixed with methanol and ultrapure water (MS grade, 1:1 v/v) to a final carbon concentration of 2.5 mg C $L^{-1}$ immediately preceding FT-ICR MS analysis. For analysis validation, an in-house DOM reference sample, collected at the Natural Energy Laboratory of Hawaii Authority in 2009 [4], was measured regularly. FT-ICR MS measurements were conducted in electrospray ionization (ESI) negative ion mode and putative chemical formulae were assigned following the same method described in Bercovici *et al.* [5].

#### Molecular characteristics of DOM

The chemical characteristics of DOM molecules were evaluated by four molecular traits, namely, H/C ratio, O/C ratio, standard Gibbs Free Energy for the half reaction of carbon oxidation (GFE) [6], and $AI_{mod}$ [7, 8]. A larger O/C ratio and lower GFE can indicate a higher degree of oxidation. GFE is associated with the oxidative degradation of organic molecules by heterotrophic organisms [6], and calculated by first estimating the nominal oxidation state of carbon (NOSC) from the equations [6]:

$$NOSC = -((-Z + 4a + b - 3c - 2d + 5e - 2f) / a) + 4 \quad (1)$$

$$GFE = 60.3 - 28.5 \times NOSC \quad (2)$$

where $a$, $b$, $c$, $d$, $e$ and $f$ are the number of atoms of elements C, H, N, O, P and S, respectively, in a given molecule, and $Z$ is the net charge of the molecule and equals zero when formula lists contained only the neutral forms of the measured negatively ionized molecular formulae. A lower H/C ratio and higher $AI_{mod}$ have been shown to correspond to a higher recalcitrance (i.e., lower bioavailability) of DOM [9, 10]. $AI_{mod}$ is calculated from the equation [7, 8]:

$$AI_{mod} = \frac{1 + C - \frac{1}{2}O - S - \frac{1}{2}(N + P + H)}{C - \frac{1}{2}O - N - S - P} \quad (3)$$

**Statistical analyses**

**Geographical patterns of compositional-level thermal responses**

The latitudinal or temperature patterns of $iCER_{avg}$ and $iCER_{var}$ are likely little affected by DOM extraction efficiency or instrument conditions. All DOM extractions followed the standard method of Dittmar *et al.* (2008) [1], and FT-ICR MS analyses were performed under consistent instrument settings. Furthermore, linear mixed-effects models were employed to assess how $iCER_{avg}$ and $iCER_{var}$ varied as a function of latitudes or water temperature when incorporating ocean or cruise identity as random effects to account for potential biases related to the order that samples were analyzed by FT-ICR MS (Table S3). These analyses confirmed the consistent patterns along latitudinal and temperature gradients as indicated in Fig. 4.

To compare the results between the surface and deep ocean in the analyses of latitudinal or temperature patterns of $iCER_{avg}$ and $iCER_{var}$, we used representative depths of ≤ 200 m and > 1,000 m, respectively. In addition, we examined the geographical patterns across finer depth intervals, namely, 0-20, 20–40, 40–60, 60–100, 100–200, 200–300, 300–1,000, 1,000–2,000, 2,000–3,000, and > 3,000 m. The results were consistent with those observed when depths were grouped into ≤ 200 m and > 1,000 m (Fig. 4), which reinforces the evidence that our findings are unlikely to be influenced by variation in depth coverage.

Furthermore, we provided additional support to compare the results between the surface and deep ocean based on water masses (Fig. S11). We confirmed that the results were robust to the two classification methods. Specifically, samples were grouped into surface and deep ocean based on water masses following the work from Schmitz [11] and Talley [12, 13] for the open ocean basins and Orsi *et al.* [14] for the Southern Ocean and Antarctic water masses. Surface water masses include Subantarctic/Subarctic Surface Water (SASW; latitude > 40 N or S and latitude < 50 S, and $\sigma_\theta < 27$ kg m$^{-3}$), Subtropical Surface Water (SSW; 20 < latitude < 40 N or S, and $\sigma_\theta < 27$ kg m$^{-3}$), Equatorial Surface Water (EqSW; latitude between 20 N and 20 S, and $\sigma_\theta < 27$ kg m$^{-3}$), and Antarctic Surface Water (AASW; latitude > 50 S, and salinity < 34.4 and depth < 250 m). Deep water masses include North Atlantic Deep Water (NADW; $\sigma_\theta > 27.5$ kg m$^{-3}$, and $\theta > 2$ ºC), Antarctic Bottom Water (AABW; $\theta < 2$ ºC, and salinity > 34.5), Circumpolar Deep Water (CDW; $\theta < 2$ ºC, and salinity > 34.5), and Pacific Deep Water (PDW; 0 < latitude < 40 N, and 100 < AOU < 250 μM).

**Environmental factors and ecosystem processes underlying compositional-level thermal responses**

We considered two key ecosystem processes—microbial transformations and photodegradation—which were quantified using the indices of $I_{bio}$ and $I_{photo}$, respectively [2]. The $I_{bio}$ was developed from the relative abundance of marker molecules of DOM derived from a three-year mesocosm experiment on DOM production by a natural microbial community of phyto- and bacterioplankton [2, 15]. The biological marker molecules are selected based on their intensities at least 30% higher than that in the reference sample that is considered refractory and stable for long time scales [2]. These molecules are also unsusceptible to photodegradation [2]. $I_{bio}$ captures

DOM molecules that are photosynthetically produced and microbially modified. Higher $I_{bio}$ indicates that microbial reworking has predominantly shaped the current DOM composition of a sample [2]. The $I_{photo}$ was developed from the relative abundance of marker molecules of DOM identified in a photodegradation experiment [2, 16]. The photochemical marker molecules were selected based on having at least 30% lower intensities after 28 days of irradiation under a solar simulator [2]. These molecules are also resistant to immediate microbial transformation [2]. $I_{photo}$ captures the DOM molecules photodegraded in the photic zone and those subsequently exported to deeper depths via physical mechanisms. Lower $I_{photo}$ indicates that photodegradation has predominantly shaped the DOM composition of a sample [2]. We note that the peaks selected for the two indices were required to have an intensity > 5% of the peak with the highest intensity in the respective sample, which ensures broad detectability across diverse environmental samples. The same set of marker peaks originally identified by Bercovici *et al.* [2] was then used directly to calculate both indices for all samples in our global ocean dataset.

We assessed the environmental factors and ecosystem processes underlying $iCER_{avg}$ and $iCER_{var}$ using three statistical methods: random forest analysis, Pearson correlation, and structural equation modelling (SEM).

First, random forest analysis was conducted to quantify the variation in $iCER_{avg}$ and $iCER_{var}$ explained by environmental factors. Strongly correlated predictors with Pearson correlations higher than 0.75 were dereplicated to reduce multicollinearity before analysis. The importance of each predictor variable was determined by evaluating the decrease in prediction accuracy (that is, increase in the mean square error between observations and out-of-bag predictions) when the data for that predictor were randomly permuted. The accuracy importance measure was estimated for each tree and averaged over the forest (2,000 trees). The relative importance of individual predictors was calculated by normalizing with the importance of the most important predictor and then multiplying by the corresponding model performance (that is, $R^2$). This analysis was conducted using the R package caret V6.0.94.

Second, two-sided Pearson correlation analysis was used to explore the relationships between $iCER_{avg}$ (or $iCER_{var}$) and the two ecosystem processes.

Third, SEM analysis was used to explore how environmental factors influenced $iCER_{avg}$ and $iCER_{var}$ directly and indirectly through the two ecosystem processes. Before modelling, all variables in the SEMs were Z-score transformed and the strong correlated predictors were dereplicated if their Pearson correlation is higher than 0.75. We used composite variables to account for the collective effects of environment factors, and the observed indicators for each composite variable were selected based on the multiple regressions for $iCER_{avg}$ and $iCER_{var}$ (Table S4). The best-fitting models were reported as that with the lowest AIC value from models with a non-statistically-significant $\chi^2$ test ($P > 0.05$), which tests whether the model structure differs from the observed data, high comparative fit index (CFI > 0.95) and low standardized root mean squared residual (SRMR < 0.05) (Table S5). We implemented the SEMs using R package lavaan V0.5.23 [17], and more details are described in previous literature [18].

**Global mapping of compositional-level thermal responses**

For the global mapping analysis, we included water depth and 12 other global environmental variables extracted from global mapping products from 1940 to 2020. Specifically, we employed the IAP gridded monthly average temperature and salinity with 1°×1° horizontal resolution at 41 vertical levels for the upper 0–2,000 m (http://www.ocean.iap.ac.cn). For radiation-related variables, we used surface latent heat flux, surface sensible heat flux, surface net long-wave (thermal) radiation, and surface net short-wave (solar) radiation from the ERA5 monthly average data with 1°×1° horizontal resolution (https://cds.climate.copernicus.eu). The environmental variables above generally exhibit inter-annual variation, while contemporary DOM compositions should be structured by historical environmental conditions. We therefore explicitly considered antecedent conditions by using the long-term (e.g., 10-years) average or standard deviation of these variables for the global mapping analysis. Practically, we aggregated dynamic covariates to annual means or standard deviations over a window size of 10 years (i.e., 10 years before the date of thermal response projecting). In summary, the 13 environmental variables used for the mapping analysis included water depth (Depth), the annual means of water temperature (Temp), salinity (Salinity), surface latent heat flux (SLHF), surface sensible heat flux (SSHF), surface net long-wave (thermal) radiation (STR), and surface net short-wave (solar) radiation (SSR), as well as the annual standard deviations of these variables: water temperature (Temp.var), salinity (Salinity.var), surface latent heat flux (SLHF.var), surface sensible heat flux (SSHF.var), surface net long-wave (thermal) radiation (STR.var), and surface net short-wave (solar) radiation (SSR.var). After reducing the multicollinearity, we included 11 environmental variables in the final models for $iCER_{avg}$ and $iCER_{var}$, as shown in Fig. 6a.

We first trained the random forest model to identify the most important variables explaining the variation in $iCER_{avg}$ and $iCER_{var}$. To fit the model, a tenfold cross-validation was conducted. Specifically, 80% of data were used to train the random forest model, and the remaining 20% to validate the model. The model performance was assessed by the root mean square error (RMSE) and coefficient of determination ($R^2$), and the best model was selected with the highest $R^2$. The relative importance of each predictor variable was determined as previously described (see statistical analysis of "random forest"). The random forest model with the lowest RMSE was then applied to predict $iCER_{avg}$ (or $iCER_{var}$) in each 1° × 1° pixel across the global ocean. The uncertainty of thermal responses was estimated as the standard error of individual predictions of 2,000 trees in the random forest model (Fig. S13). The mapping analysis was performed at 39 water depths for the upper 0–2,000 m in 1950 and 2020.

We further examined the temporal changes in $iCER_{avg}$ and $iCER_{var}$ from 1950 to 2020 in each 1° × 1° pixel across the global ocean. Because iCER values can be either positive or negative and may change sign between 1950 and 2020, a single formula cannot robustly quantify percentage change in all scenarios. To ensure numerical stability and comparability, we calculated percentage changes using three alternative methods (Methods 1–3), each of which applies a specific equation depending on the sign and relative magnitude of $\mathrm{iCER}_{2020}$ and $\mathrm{iCER}_{1950}$. The equations used in each method are provided as follows:

$$\text{Percentage change} = \frac{\mathrm{iCER}_{2020} - \mathrm{iCER}_{1950}}{|\mathrm{iCER}_{1950}|} \times 100 \tag{4}$$

$$\text{Percentage change} = \left(\frac{\mathrm{iCER}_{2020}}{\mathrm{iCER}_{1950}} - 1\right) \times 100 \tag{5}$$

$$\text{Percentage change} = \left(\frac{\mathrm{iCER}_{2020}}{|\mathrm{iCER}_{1950}|} + 1\right) \times 100 \tag{6}$$

$$\text{Percentage change} = -\frac{|-|\mathrm{iCER}_{2020}| - |\mathrm{iCER}_{1950}||}{|\mathrm{iCER}_{1950}|} \times 100 \tag{7}$$

$$\text{Percentage change} = \left|\frac{|\mathrm{iCER}_{2020}|}{|\mathrm{iCER}_{1950}|} - 1\right| \times 100 \tag{8}$$

$$\text{Percentage change} = -\left|\frac{|\mathrm{iCER}_{2020}|}{|\mathrm{iCER}_{1950}|} - 1\right| \times 100 \tag{9}$$

$$\text{Percentage change} = \sqrt{\frac{(\mathrm{iCER}_{2020} - \mathrm{iCER}_{1950})^2}{(\mathrm{iCER}_{1950})^2}} \times 100 \tag{10}$$

$$\text{Percentage change} = -\sqrt{\frac{(\mathrm{iCER}_{2020} - \mathrm{iCER}_{1950})^2}{(\mathrm{iCER}_{1950})^2}} \times 100 \tag{11}$$

where $\mathrm{iCER}_{2020}$ and $\mathrm{iCER}_{1950}$ are the $\mathrm{iCER}_{avg}$ (or $\mathrm{iCER}_{var}$) in 2020 and 1950, respectively. The conditions under which each equation is applied and the corresponding rationale are summarized in Table S6. All three methods produced the same results. Statistical significance of the temporal changes between 1950 and 2020 was tested with Student's t-test. To visualize the depth profiles of temporal changes in thermal responses, the vertical levels were interpolated into higher vertical resolution (1-m interval) based on linear interpolation.

**Supplementary Tables**

**Table S1**. Summary of study aims and the corresponding statistical analyses. The locations indicate the figures in which these analyses were employed.

| **Overall aims** | | **Statistical analyses and their aims** | | **Locations** |
|---|---|---|---|---|
| **Aim 1:** MER-trait associations | Assessing how the molecular composition and traits (e.g., recalcitrance and oxidation state) of DOM changed towards stronger thermal responses of individual molecules | Moving-window analysis | • To examine the continuous and/or sharp transitions in molecular composition or traits along a MER continuum<br>• A MER continuum from negative to positive was generated by categorizing all molecules into 390 equal-sized bins (200 molecules per bin), with each bin representing a DOM constituent. | Fig. 2 |
| **Aim 1:** Molecule-level persistence | Assessing the persistence of the DOM molecules with negative or positive MERs throughout the water column | Calculation of apparent carbon concentrations of DOM molecules | • To be quantified as the normalized mass peak intensities weighted by the respective SPE-DOC concentration<br>• Calculated for each molecule individually or each DOM constituent (i.e., the molecules in each of the 390 molecule bins) | Figs. 3, S2 |
| | | Spearman correlation | • To analyze depth profiles of apparent carbon concentrations of DOM molecules at each station<br>• Persistence of DOM molecules was indicated by statistically non-significant depth profiles of apparent carbon concentrations of DOM molecules<br>• Percentages of statistically non-significant depth profiles were calculated for individual molecules or the molecule bins across the stations | Figs. 3, S2 |
| **Aim 2:** Geographical patterns | Exploring patterns of compositional-level thermal responses along the gradients of latitude, water depth, | Linear regression | • To analyze the relationships between $iCER_{avg}$ (or $iCER_{var}$) and latitude (or water depth and temperature) | Figs. 4, S3, S11 |

| | | | | |
|---|---|---|---|---|
| | and water temperature | | | |
| **Aim 3:** iCER-DOC associations | Examining how compositional-level thermal responses could predict recalcitrant DOC concentrations | Linear regression | • To analyze the relationships between $iCER_{avg}$ (or $iCER_{var}$) and the apparent carbon concentrations of DOM constituents across a MER continuum from negative to positive, corresponding to a gradient from more recalcitrant to more labile molecules<br>• The apparent carbon concentrations of DOM constituents across a MER continuum were defined as the summed apparent carbon concentrations of DOM molecules in each of the 390 molecule bins | Figs. 5b, d |
| **Aim 3:** Underlying processes | Assessing the environmental factors and ecosystem processes underlying compositional-level thermal responses | Random forest analysis | • To quantify the variation in $iCER_{avg}$ and $iCER_{var}$ explained by environmental factors | Fig. S4 |
| | | Pearson correlation | • To explore the relationships between $iCER_{avg}$ (or $iCER_{var}$) and the two ecosystem processes<br>• The two ecosystem processes of microbial reworking and photochemical degradation were quantified using the indices of $I_{bio}$ and $I_{photo}$, respectively | Figs. 5e, g |
| | | Structural equation modelling (SEM) analysis | • To explore how environmental factors influenced $iCER_{avg}$ and $iCER_{var}$ directly and indirectly through $I_{bio}$ and $I_{photo}$ | Figs. 5f, h |
| **Implication:** iCER projection & implication | Estimating the spatiotemporal changes in compositional-level thermal responses across the global ocean between 1950 and 2020 | Machine learning-based random forest model | • To identify the most important variables explaining the variation in $iCER_{avg}$ and $iCER_{var}$<br>• To predict $iCER_{avg}$ and $iCER_{var}$ in each 1° × 1° pixel across the global ocean at 39 water depths for the upper 0–2,000 m in 1950 and 2020 | Figs. 6a-c, S7-8 |
| | | Percentage change analysis | • To examine the temporal changes in $iCER_{avg}$ and $iCER_{var}$ between 1950 and 2020 | Figs. 6d, S9 |
| | Estimating the impact of changes in $iCER_{var}$ | Linear regression | • To calculate the linear slope of the relationship between the | Table S2 |

| | | | | |
|---|---|---|---|---|
| | from 1950 to 2020 on the concentration changes of recalcitrant DOC in the global ocean | | $iCER_{var}$ and the apparent carbon concentration of molecules with negative MERs<br>• The concentration of recalcitrant DOC was defined as the apparent carbon concentrations of molecules with negative MERs (Spearman $\rho < 0$; $P \leq 0.05$) and divided by DOM extraction efficiency | |
| | | Estimation of changes in recalcitrant DOC | • To calculate the changes in the concentration of recalcitrant DOC based on the linear slope described above using Equation 5 (see Methods) | Table S2 |

**Table S2**. The changes in recalcitrant DOC (Tg C $yr^{-1}$) for each additional increase in the diversity of thermal responses ($iCER_{var}$) from 1950 to 2020. Recalcitrant DOC was calculated as the apparent carbon concentrations of molecules with negative MERs and divided by DOM extraction efficiency. The volumes of seawater at different depths are available from Costello *et al.* [19].

| Water depth (m) | Seawater volume ($km^3$) | $iCER_{var}$ changes (%) [#] | Recalcitrant DOC changes (Tg C $yr^{-1}$) [#] | Formula of relationships between recalcitrant DOC and $iCER_{var}$ |
|---|---|---|---|---|
| < 100 | 796,975 | - 0.039 (±0.0244) | - 0.00417 (±0.00262) | Recalcitrant DOC = 101.663 × $iCER_{var}$ – 32.94 |
| 101-500 | 3,238,442 | - 0.002 (±0.0138) | - 0.00940 (±0.07888) | Recalcitrant DOC = 134.476 × $iCER_{var}$ – 45.68 |
| > 1,000 | $1.3 \times 10^9$ | + 0.047 (±0.0159) | + 9.87 (±3.35) | Recalcitrant DOC = 125.434 × $iCER_{var}$ – 41.64 |

[#] $iCER_{var}$ changes and recalcitrant DOC changes are presented as the means ± SD averaged across water depths.

**Table S3**. Summary of the model fit statistics for linear mixed-effects models. We assessed the strength ($iCER_{avg}$) and diversity ($iCER_{var}$) of DOM thermal responses as a function of absolute latitude or water temperature after accounting for ocean or cruise identity as random effects.

| | | $iCER_{avg}$ | | | $iCER_{var}$ | | |
|---|---|---|---|---|---|---|---|
| Fixed factor | Random factor | Slope | $R^2$ | $P$ | Slope | $R^2$ | $P$ |
| Absolute latitude | None | -0.0010 | 0.21 | <0.001 | -0.0001 | 0.03 | <0.001 |
| | Oceans | -0.0010 | 0.19 | <0.001 | -0.0001 | 0.06 | 0.005 |
| | Cruises | -0.0007 | 0.10 | 0.0015 | -0.0002 | 0.08 | <0.001 |
| Water temperature | None | 0.0034 | 0.54 | <0.001 | 0.0006 | 0.30 | <0.001 |
| | Oceans | 0.0034 | 0.54 | <0.001 | 0.0006 | 0.33 | <0.001 |
| | Cruises | 0.0033 | 0.48 | <0.001 | 0.0008 | 0.45 | <0.001 |

**Table S4**. Formulae to calculate composite variables for structure equation models of the compositional-level thermal responses of oceanic DOM (iCER$_{avg}$ and iCER$_{var}$) for all samples, and samples at depths of ≤ 200 m and > 1,000 m. The obtained composite variables of environments (Envi) were used in Figs. 5f and h.

| Thermal response | Water depth (m) | Formula |
|---|---|---|
| iCER$_{avg}$ | All samples | -0.09 × Depth ($\log_{10}$) + 0.74 × Temperature + -0.09 × Salinity |
| | ≤ 200 | 0.80 × Temperature + -0.11 × Salinity |
| | > 1,000 | 0.19 × Depth ($\log_{10}$) + 0.25 × Temperature + -0.13 × Salinity |
| iCER$_{var}$ | All samples | 0.16 × Latitude + -0.30 × Depth ($\log_{10}$) + 0.42 × Temperature + 0.15 × Salinity |
| | ≤ 200 | 0.29 × Temperature + 0.21 × Salinity |
| | > 1,000 | 0.42 × Latitude + 0.36 ×Temperature + 0.44 × Salinity |

**Table S5**. Summary of the model fit statistics evaluated for standardized structural equation model (SEM). We explored the potential links between predictor variables and the compositional-level thermal responses of oceanic DOM (iCER$_{avg}$ and iCER$_{var}$) for all samples, and samples at depths of ≤ 200 m and > 1,000 m. $\chi^2$: Chi-square. *P*: p-value of chi-square test. df: Degrees of freedom. CFI: Comparative fit index. SRMR: Standardized root mean square residual. AIC: Akaike information criterion.

| Thermal response | Water depth (m) | df | $\chi^2$ | *P* | CFI | SRMR | AIC |
|---|---|---|---|---|---|---|---|
| | All samples | 1 | 1.647 | 0.199 | 1.000 | 0.009 | 5993.0 |
| iCER$_{avg}$ | ≤ 200 | 1 | 1.340 | 0.247 | 1.000 | 0.012 | 4005.6 |
| | > 1,000 | 6 | 0.008 | 0.927 | 1.000 | 0.002 | 1554.6 |
| | All samples | 1 | 1.120 | 0.290 | 1.000 | 0.007 | 6191.5 |
| iCER$_{var}$ | ≤ 200 | 1 | 2.264 | 0.132 | 0.999 | 0.017 | 4195.3 |
| | > 1,000 | 1 | 2.892 | 0.089 | 0.992 | 0.058 | 1766.5 |

**Table S6**. Summary of the methods used to calculate percentage changes in $iCER_{avg}$ and $iCER_{var}$ from 1950 to 2020.

| **Method** | **Equation No.** | **Condition ($iCER_{2020}$ vs $iCER_{1950}$)** | **Rationale** |
|---|---|---|---|
| **Method 1** | Eq. 4 | General case where a simple ratio is meaningful | Standard percentage change when division by $iCER_{1950}$ does not introduce instability |
| **Method 2** | Eq. 5 | $iCER_{2020} > 0$ and $iCER_{1950} > 0$ | Both positive → ratio reliably reflects proportional change |
| **Method 2** | Eq. 6 | $iCER_{2020} > 0$ and $iCER_{1950} \leq 0$ | Sign change → add 1 to avoid division by a value crossing zero |
| **Method 2** | Eq. 7 | $iCER_{2020} \leq 0$ and $iCER_{1950} > 0$ | Sign change → absolute formulation avoids undefined or misleading ratios |
| **Method 2** | Eq. 8 | $iCER_{2020} \leq 0$ and $iCER_{1950} \leq 0$ and $iCER_{2020} > iCER_{1950}$ | Both negative and increasing → magnitude-based ratio is valid |
| **Method 2** | Eq. 9 | $iCER_{2020} \leq 0$ and $iCER_{1950} \leq 0$ and $iCER_{2020} < iCER_{1950}$ | Both negative and decreasing → sign-sensitive (negative) ratio required |
| **Method 3** | Eq. 10 | $iCER_{2020} > iCER_{1950}$ | Root-mean-square formulation captures magnitude increase regardless of sign |
| **Method 3** | Eq. 11 | $iCER_{2020} \leq iCER_{1950}$ | Root-mean-square formulation captures magnitude decrease regardless of sign |

### Supplementary Figures

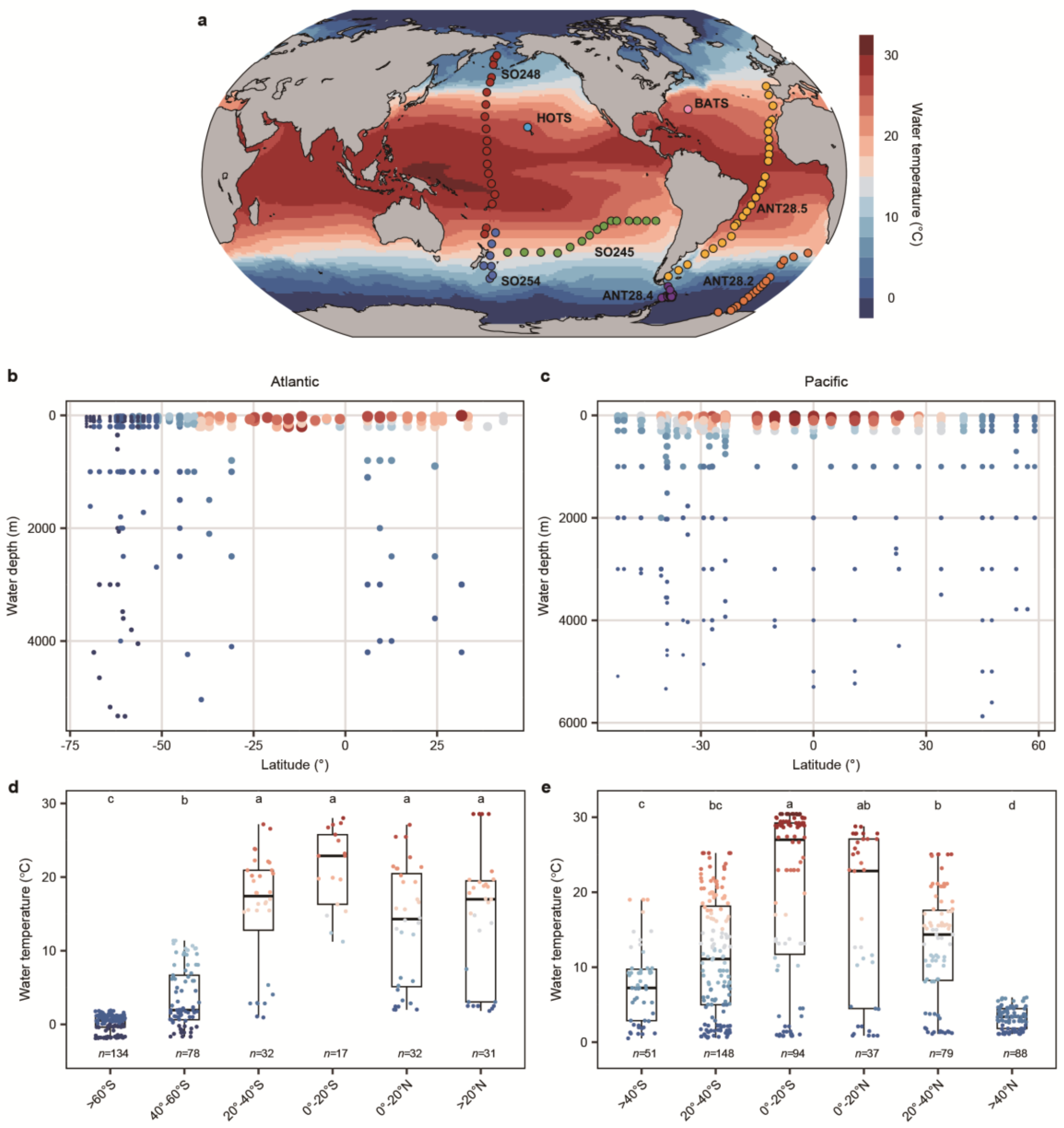


**Figure S1**. Sampling sites over depth at 124 stations during six cruises in the Atlantic, Southern, and Pacific oceans. (a) Stations visited during three *R.V. Sonne* cruises—SO245 (December 2015 to January 2016; green), SO248 (May 2016; red) and SO254 (January and February 2017; blue)—in the Pacific Ocean, and during three *R.V. Polarstern* cruises—ANT-XXVIII/2 (ANT28.2; orange), ANT-XXVIII/4 (ANT28.4; purple) and ANT-XXVIII/5 (ANT28.5; yellow)—in the Atlantic and Southern Ocean during austral spring and summer (Dec 2011 - May 2012). Stations are overlaid on global map with annual mean water temperatures at the depth of 1 m in 2020 (http://www.ocean.iap.ac.cn/). (b, c) Sampling sites covering the entire water column across depth gradients from 5 to 5,874 m at each station in the Atlantic Ocean (including Southern Ocean; b) and Pacific Ocean (c). Dot colors and sizes denote the annual mean water temperatures in 2020 at each sampling site, with higher temperatures shown by larger sized dots in darker shades of red and lower temperatures shown by smaller sized dots in darker shades of blue. (d, e) Water temperature across latitudinal zones in the Atlantic Ocean (including the Southern Ocean; d) and the Pacific Ocean (e). Dots in the boxplots represent water temperatures for individual samples. Boxes extend from 25$^{th}$ to 75$^{th}$ percentiles (first and third quartiles), with the median indicated by the central line. *n* denotes the number of samples in each latitudinal zone. Different letters indicate significant ($P \leq 0.05$) differences based on the Kruskal-Wallis test.

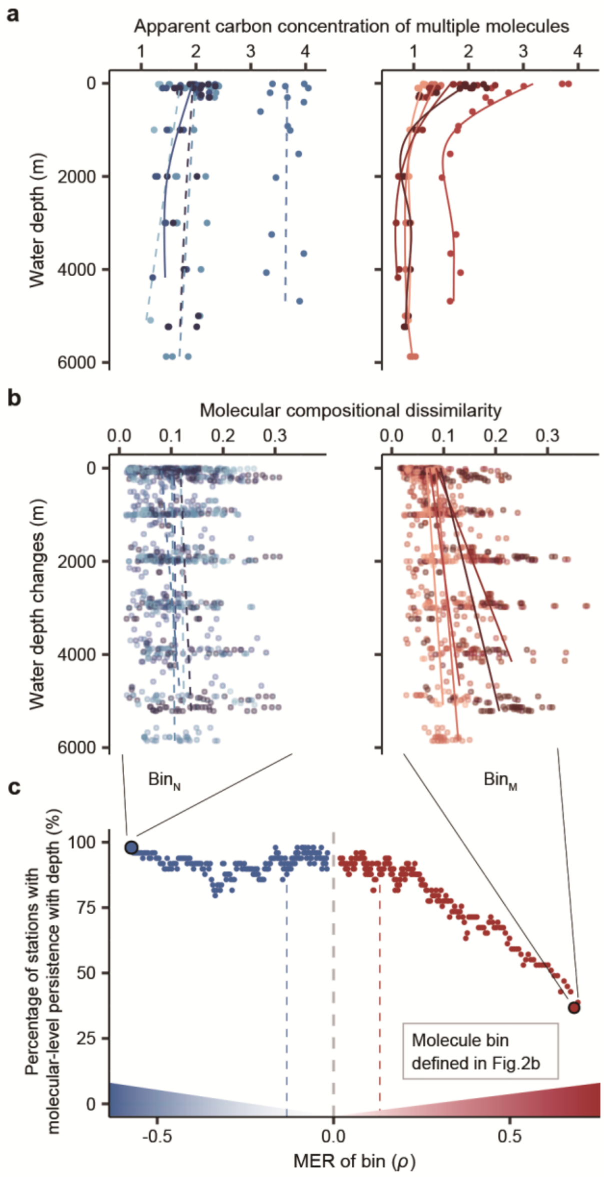


**Figure S2**. Depth profiles of the apparent carbon concentrations for DOM constituents (i.e., molecule bins) along the continuum of negative to positive molecule-specific thermal responses (MERs) of oceanic DOM. Empirical apparent carbon concentration-depth profiles (a, b) are illustrated for two example bins with negative (blue) and positive (red) MERs at five stations. We considered the relationships between the apparent carbon concentrations of multiple molecules summed in each bin and the water depth (a), and the relationships between its compositional dissimilarity (Bray-Curtis) and the differences in depth (b). At each station, molecular-level persistence was reflected by the statistically non-significant relationships between compositional dissimilarity and depth changes. The percentages of the statistically non-significant relationships across stations were shown along the MER continuum in (c). The depth profiles of apparent carbon concentrations in (a) are visualized with generalized additive models with a k of 5. Statistical significance of the relationships in (b) and (c) was tested with a Mantel test with 999 permutations. Blue and red dashed lines in (c) highlight the cutoff of 100% statistically significantly negative and positive MERs for molecules in each bin, respectively.

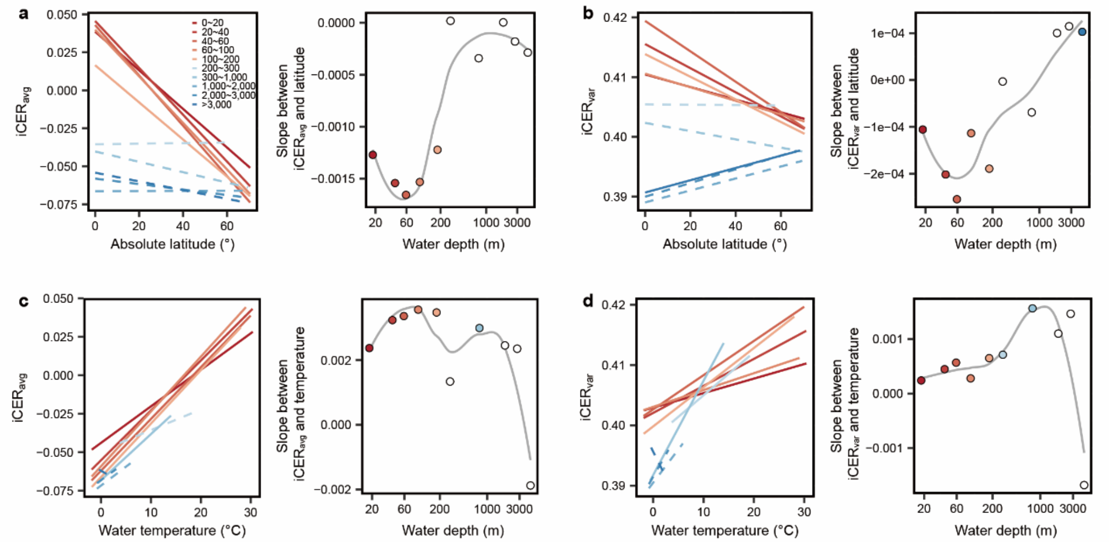


**Figure S3**. The compositional-level thermal responses of oceanic DOM along latitudinal and temperature gradients. We plotted the strength (iCER$_{avg}$; a, c) and diversity (iCER$_{var}$; b, d) of thermal responses against absolute latitude (a, b) and water temperature (c, d) across finer depth intervals than those in Fig. 4, specifically 0-20, 20-40, 40-60, 60-100, 100-200, 200-300, 300-1,000, 1,000-2,000, 2,000-3,000, and > 3,000 m. Statistical significance of linear model fits with one-sided F-statistics is indicated by solid ($P \leq 0.05$) or dotted ($P > 0.05$) lines. The linear slopes of the relationships for each depth group were further plotted against their mean depths (right panels). Solid and open circles indicate the statistically significant ($P \leq 0.05$) and non-significant ($P > 0.05$) linear slopes, respectively. The depth profiles of linear slopes were visualized with loess regression models (grey lines). These latitudinal or temperature patterns were consistent with those observed when depths were grouped into ≤ 200 m and > 1,000 m (Fig. 4), indicating that our findings are unlikely to be influenced by variation in depth coverage.

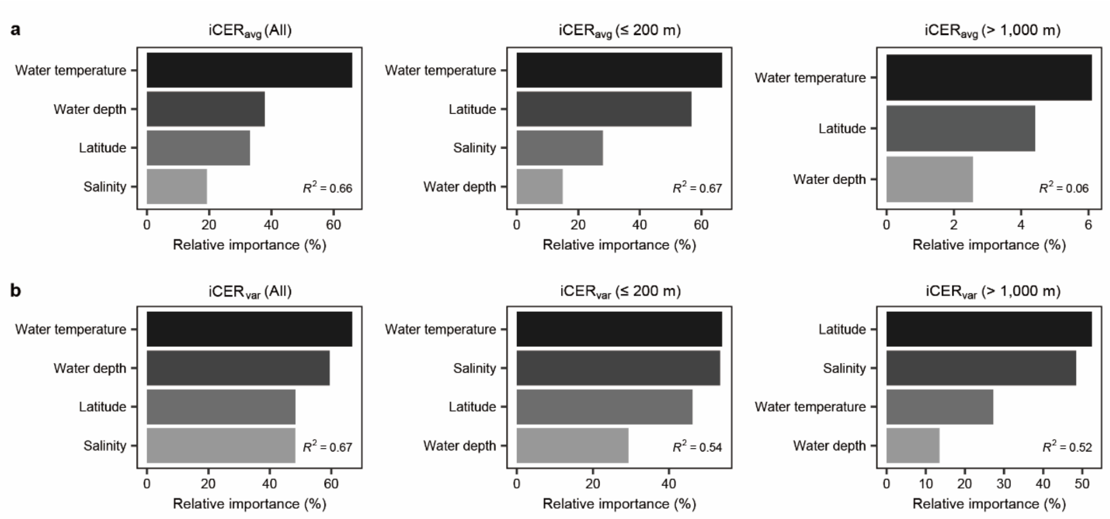


**Figure S4**. The relative importance of explanatory environmental variables on the compositional-level thermal responses of oceanic DOM. We illustrated the relative importance of environmental variables on iCER$_{avg}$ (a) and iCER$_{var}$ (b), determined by random forest analyses, for all DOM samples (left panels), and the samples at depths of ≤ 200 m (middle panels) and > 1,000 m (right panels). The adjusted $R^2$ are the total explained variances. The relative importance (%) of each variable for iCER$_{avg}$ and iCER$_{var}$ are shown as bar plots.

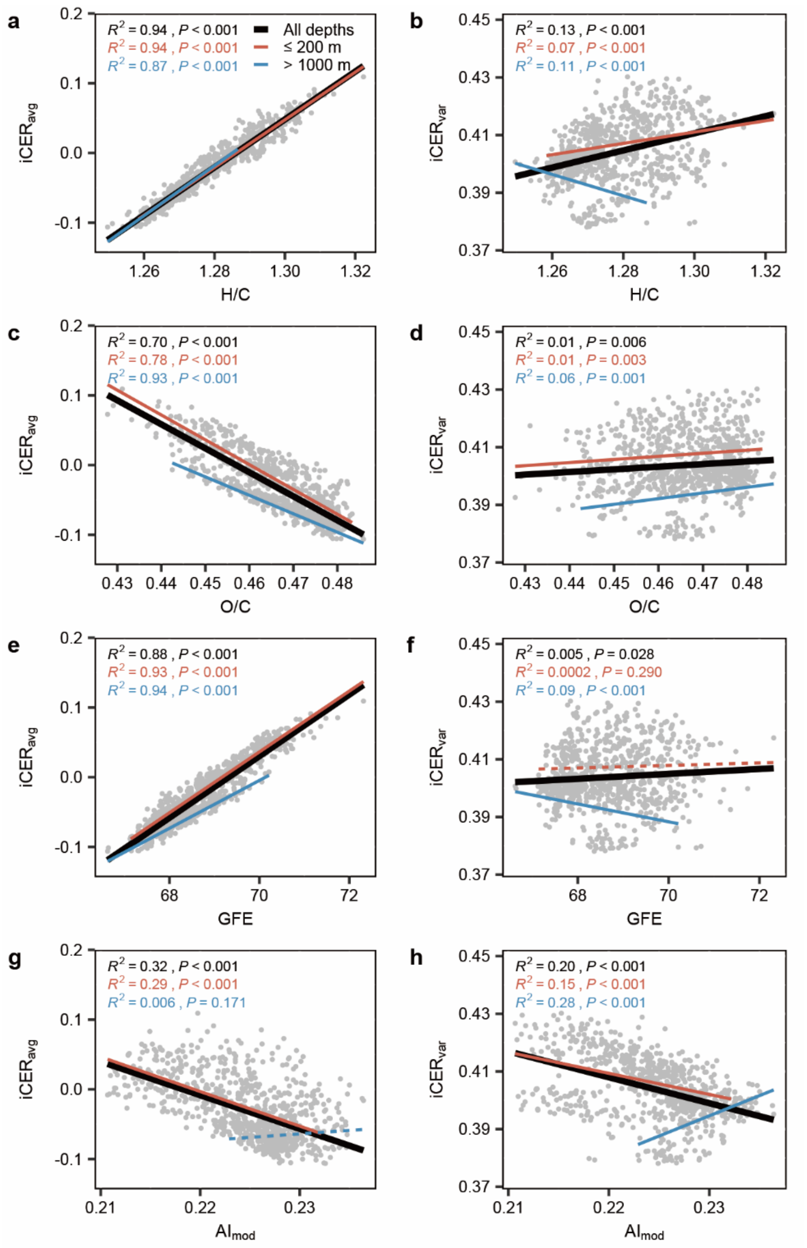


**Figure S5**. The relationships between the compositional-level thermal responses of oceanic DOM and the compositional-level DOM traits. We plotted the strength (iCER$_{avg}$; a, c, e, g) and diversity (iCER$_{var}$; b, d, f, h) of thermal responses against H/C ratio (a, b), O/C ratio (c, d), Gibbs free energy (GFE; e, f), and the modified aromaticity index (AI$_{mod}$; g, h) for all DOM samples (black lines), and the samples at depths of ≤ 200 m (red lines) and > 1,000 m (blue lines). Statistical significance of linear model fits with one-sided F-statistics is indicated by solid ($P \leq 0.05$) or dotted ($P > 0.05$) lines.

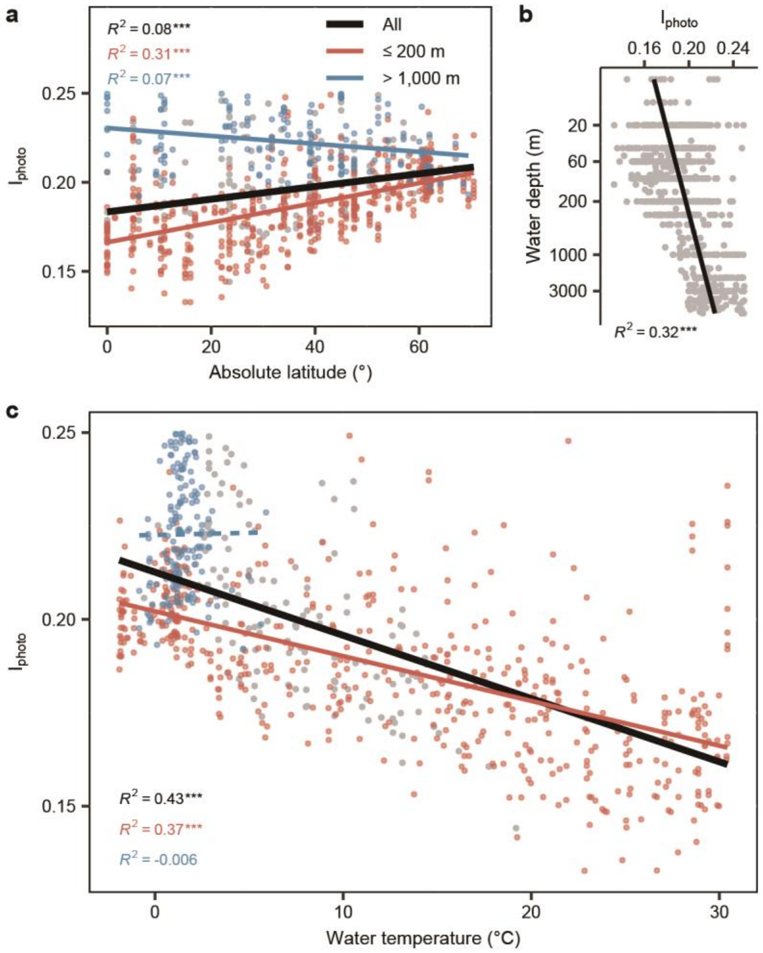


**Figure S6**. The photochemical degradation index ($I_{photo}$) of oceanic DOM along geographic and temperature gradients. We plotted $I_{photo}$ against absolute latitude (a), water depth (b), and water temperature (c) for all DOM samples (black lines, $n$ = 821). For latitude and temperature, we further considered samples in two depth categories of ≤ 200 m (red lines, $n$ = 523) and > 1,000 m (blue lines, $n$ = 159). Statistical significance of linear model fits with one-sided F-statistics is indicated by solid ($P \leq$ 0.05) or dotted ($P$ > 0.05) lines. To evaluate heteroscedasticity in $I_{photo}$-temperature relationships, we applied the Breusch-Pagan test and report only the weighted linear regression outputs for these models, with statistical significance indicated by solid ($P$ ≤ 0.05) or dotted ($P$ > 0.05) lines.

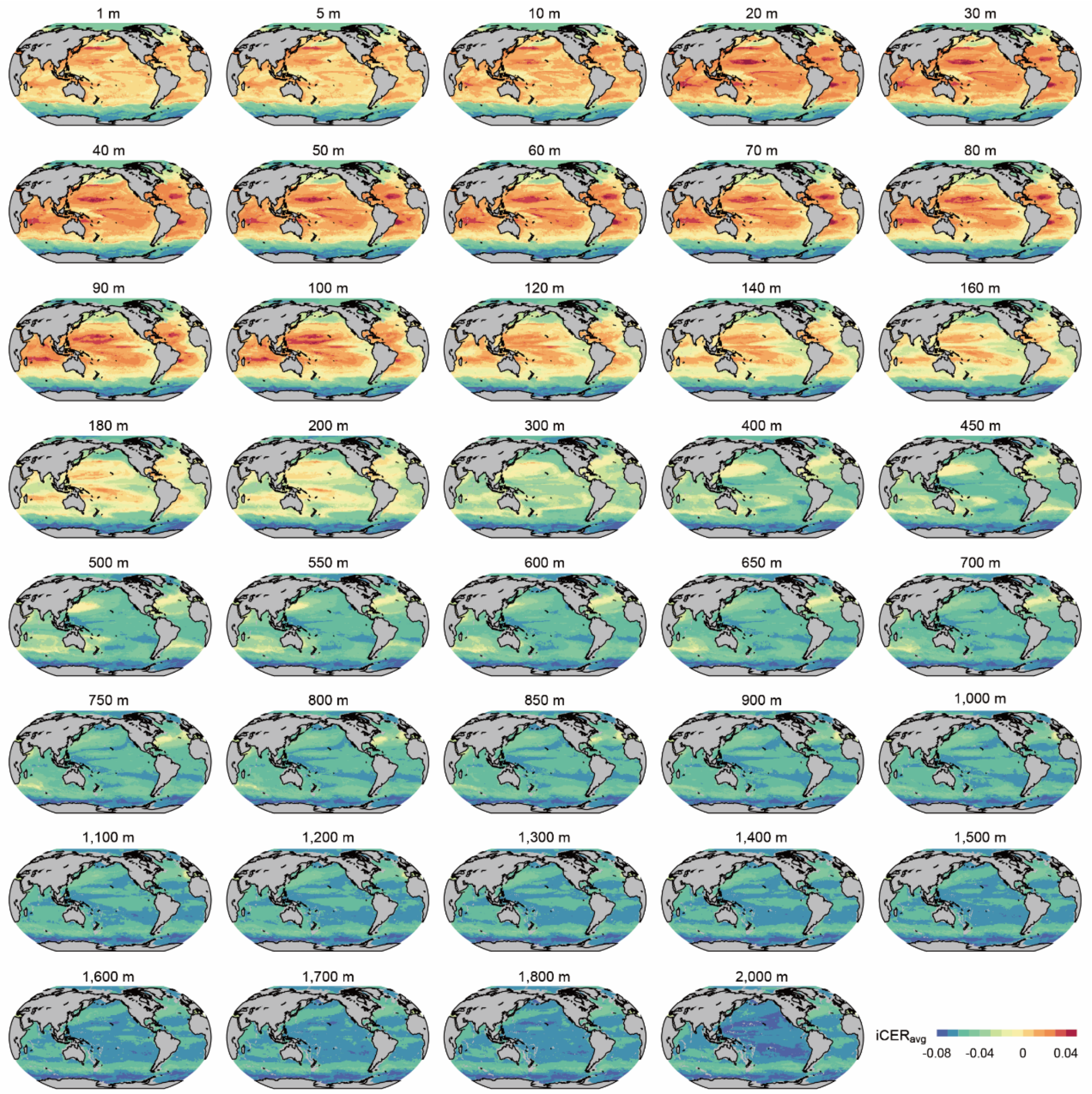


**Figure S7**. Global variations in the strength of compositional-level thermal responses of oceanic DOM (iCER$_{avg}$) in 2020 from the surface to 2,000 m. The maps show the modelled iCER$_{avg}$ across the globe, and the maps have a spatial resolution of 1°.

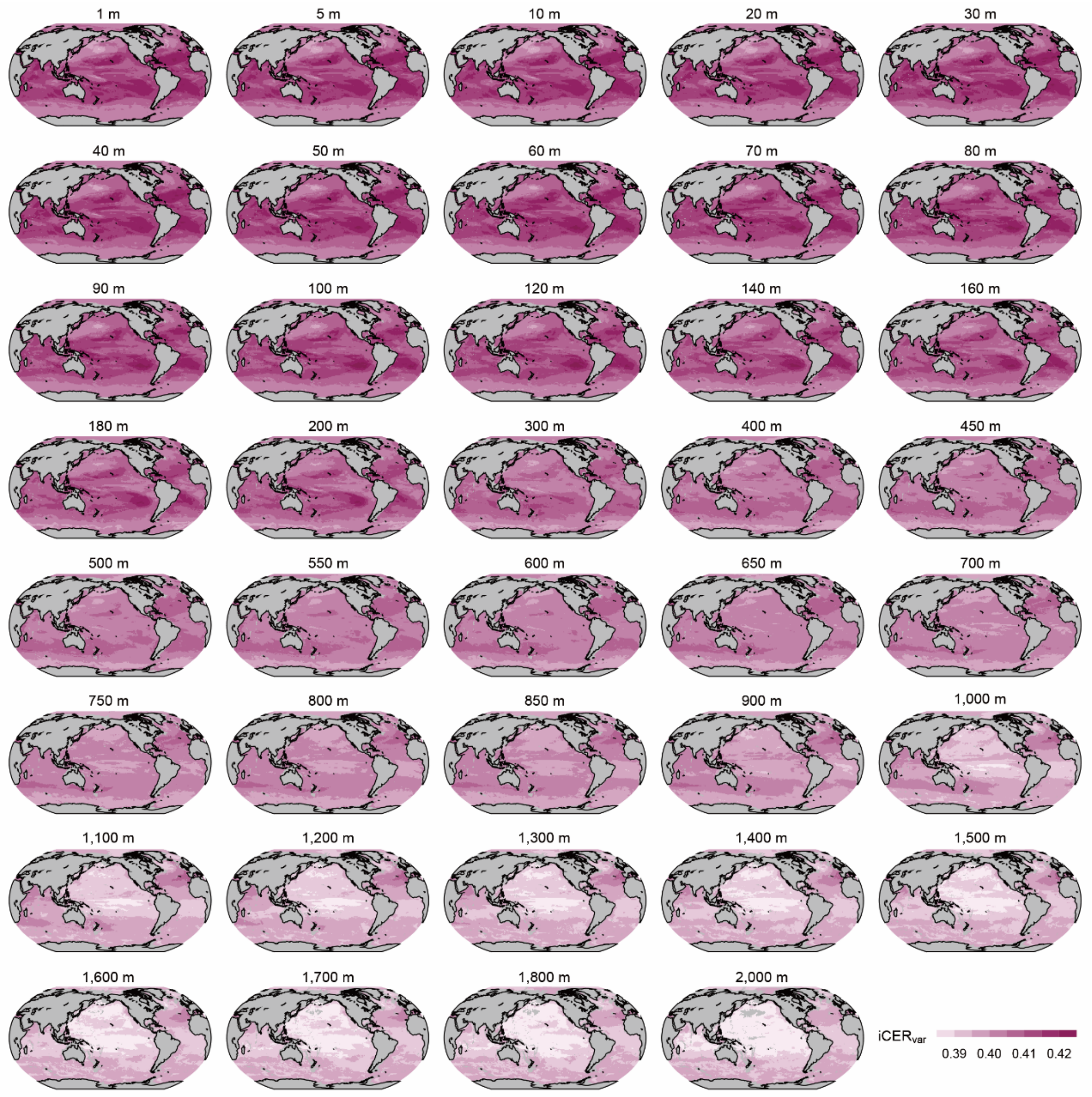


**Figure S8**. Global variations in the diversity of compositional-level thermal responses of oceanic DOM (iCER$_{var}$) in 2020 from the surface to 2,000 m. The maps show the modelled iCER$_{var}$ across the globe, and the maps have a spatial resolution of 1°.

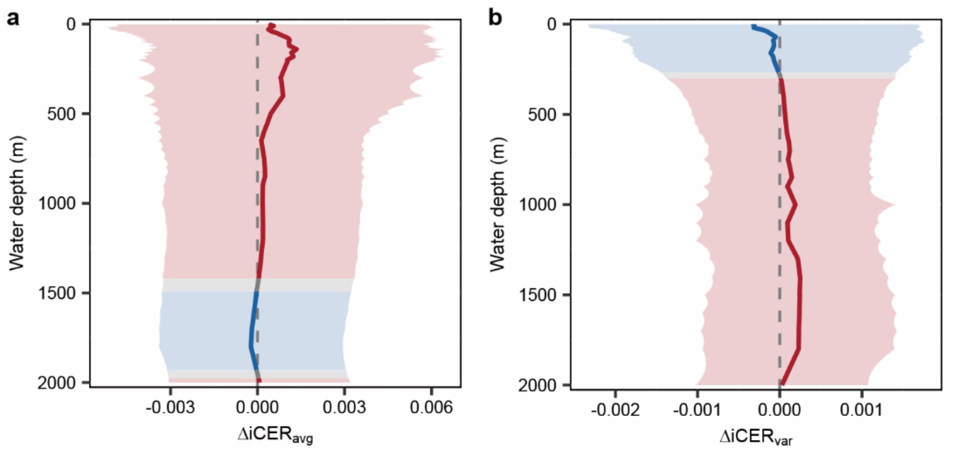


**Figure S9**. The depth profiles of temporal changes in global mean iCER$_{avg}$ (a) and iCER$_{var}$ (b) between 1950 and 2020 from the surface to 2,000 m. Blue and red line segments indicate the decreased and increased, respectively, changes in the iCER$_{avg}$ or iCER$_{var}$, and the shadings denote their standard deviations. Statistical significance of the temporal changes was tested with Student's *t*-test, with red/blue line segments indicating significant changes ($P \leq 0.05$) and grey line segments indicating non-significant changes ($P > 0.05$).

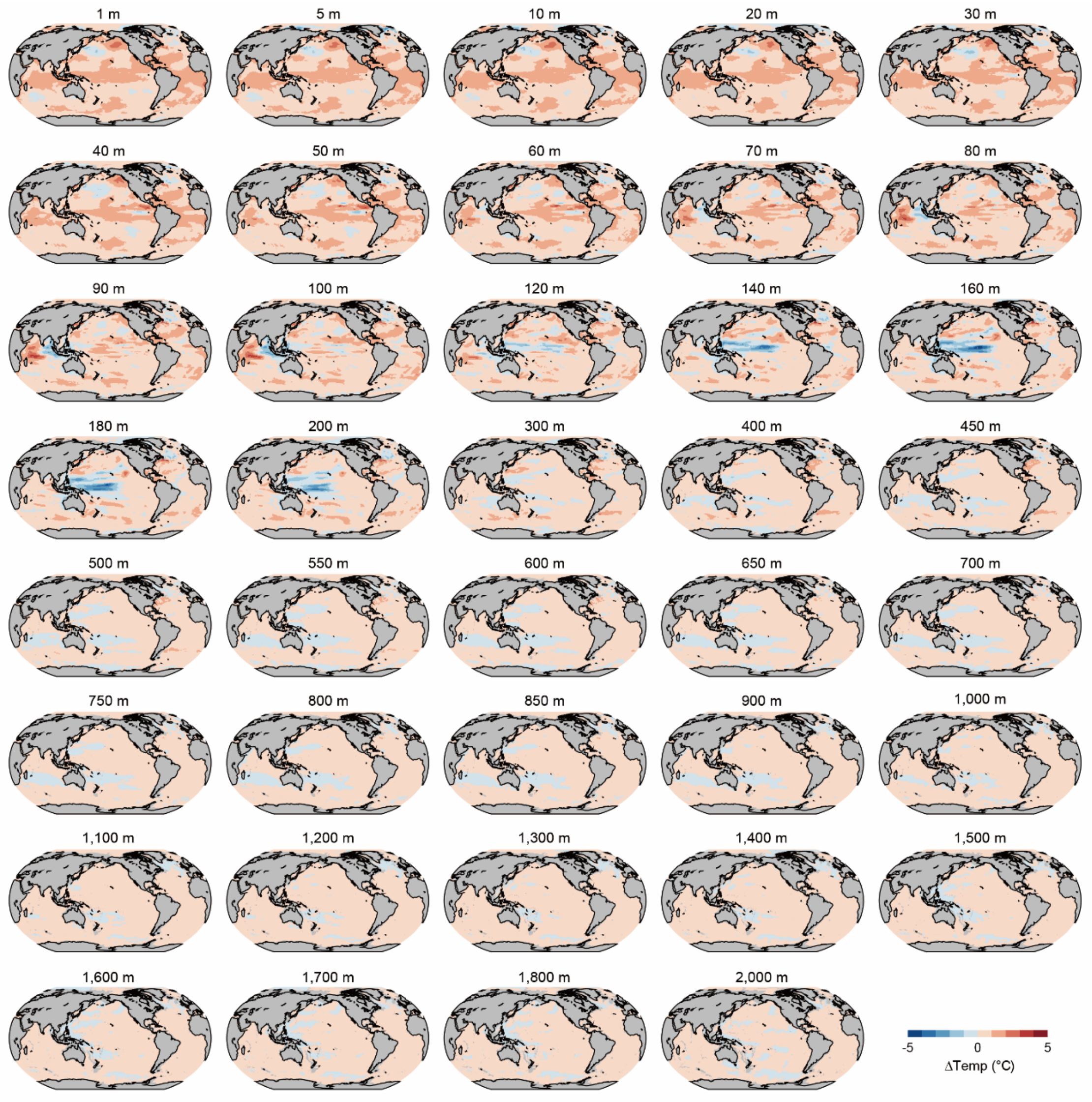


**Figure S10**. Changes in water temperatures between 1950 and 2020 from the surface to 2,000 m. The maps show the spatial distribution of water temperature across the globe, and the maps have a spatial resolution of 1°.

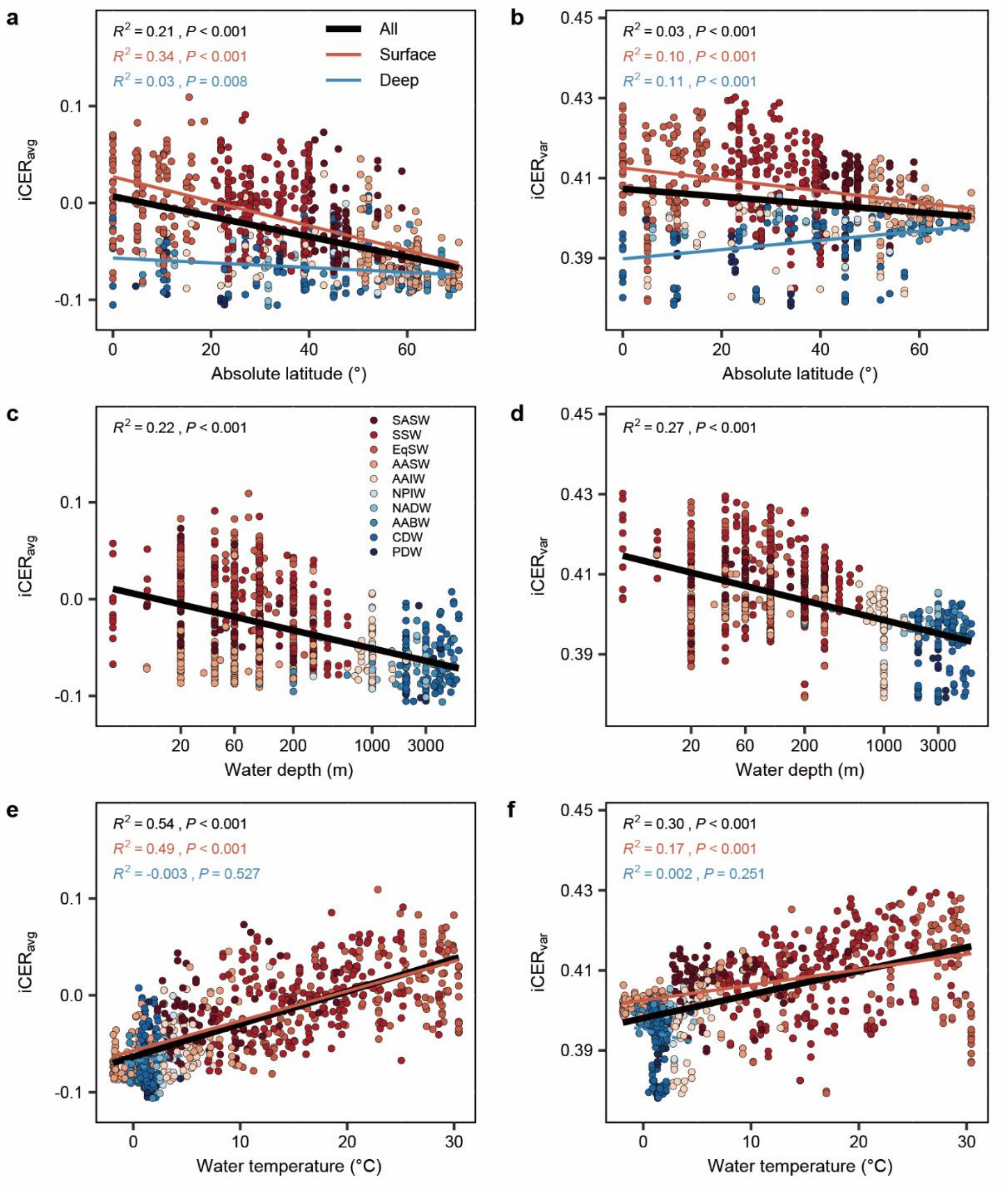


**Figure S11**. The compositional-level thermal responses of oceanic DOM along geographical and temperature gradients. We plotted the strength (iCER$_{avg}$; a, c, e) and diversity (iCER$_{var}$; b, d, f) of thermal responses against absolute latitude (a, b), water depth (c, d), and water temperature (e, f) for all DOM samples (black lines). For the latitude and temperature, we further considered two depth categories, that is surface (red lines) and deep (blue lines) waters. These depth categories are based on water masses, which are defined in Supplementary Methods. Statistical significance of linear model fits with one-sided F-statistics is indicated by solid ($P \leq 0.05$) or dotted ($P > 0.05$) lines.

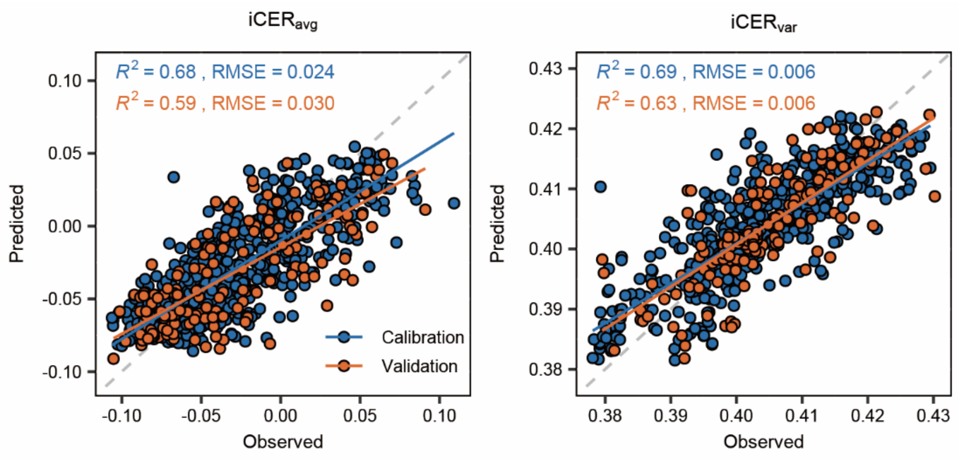


**Figure S12**. The performance of machine learning based random forest models for assessing the influences of *in situ* measured environmental variables on the compositional-level thermal responses of oceanic DOM. The environmental variables included water temperature, salinity, water depth, and latitude. The root mean square error (RMSE) and coefficient of determination ($R^2$) are shown for $iCER_{avg}$ (a) and $iCER_{var}$ (b).

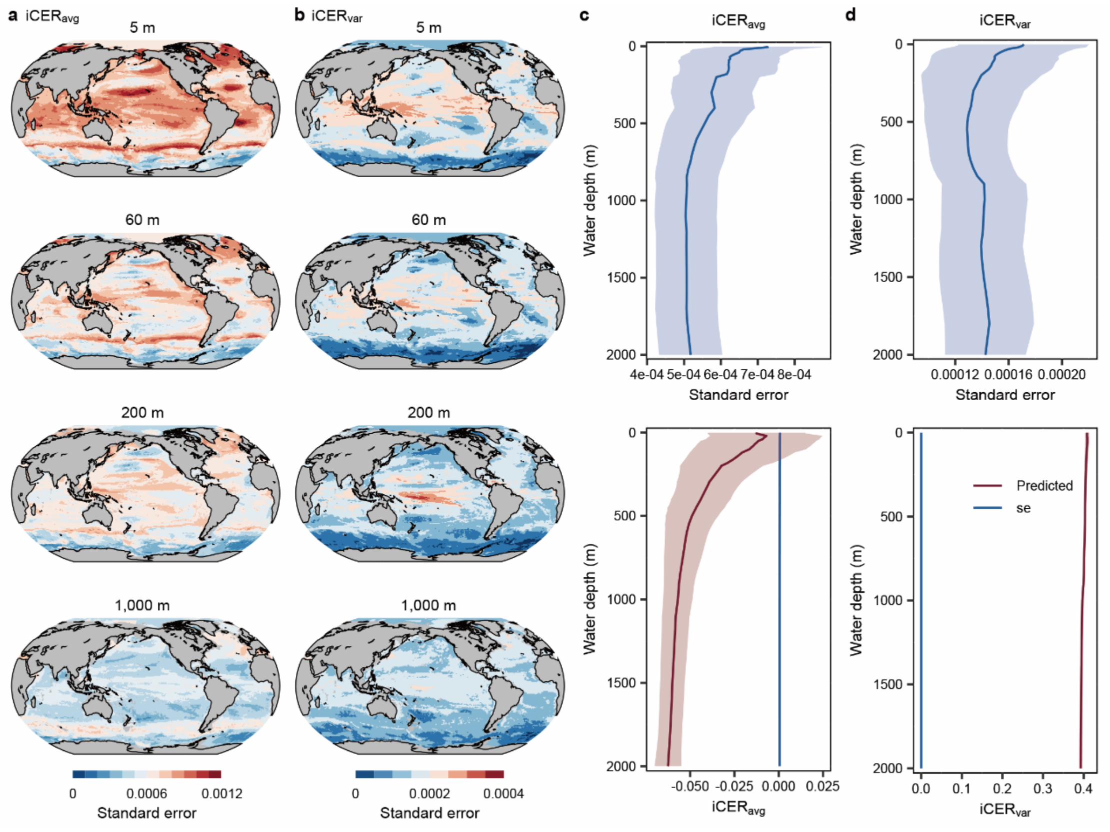


**Figure S13**. Uncertainties in the modelled compositional-level thermal responses of oceanic DOM in 2020 from the surface to 2,000 m. The uncertainties were estimated as the standard error of individual predictions of 2,000 trees in the machine learning based random forest models. (a, b) Maps of standard error for $iCER_{avg}$ (a) and $iCER_{var}$ (b) across the globe. (c, d) The mean standard error (blue lines) across the latitudinal gradient. The red line in the lower panels is the modelled $iCER_{avg}$ or $iCER_{var}$. The shadings denote their standard deviations.
